# SAMHD1 limits the efficacy of forodesine in leukaemia by protecting cells against cytotoxicity of dGTP

**DOI:** 10.1101/2020.02.17.951517

**Authors:** Tamara Davenne, Jenny Klintman, Sushma Sharma, Rachel E. Rigby, Chiara Cursi, Anne Bridgeman, Bernadeta Dadonaite, Kim De Keersmaecker, Peter Hillmen, Andrei Chabes, Anna Schuh, Jan Rehwinkel

## Abstract

The anti-leukaemia agent forodesine causes cytotoxic overload of intracellular deoxyguanosine triphosphate (dGTP) but is efficacious only in a subset of patients. We report that SAMHD1, a phosphohydrolase degrading deoxyribonucleoside triphosphates (dNTPs), protected cells against the effects of dNTP imbalances. SAMHD1-deficient cells induced intrinsic apoptosis upon provision of deoxyribonucleosides, particularly deoxyguanosine (dG). Moreover, dG and forodesine acted synergistically to kill cells lacking SAMHD1. Using mass cytometry, we found that these compounds killed SAMHD1-deficient malignant cells from patients with chronic lymphocytic leukaemia (CLL). Normal cells and CLL cells from patients without *SAMHD1* mutation were unaffected. We therefore propose to use forodesine as a precision medicine for leukaemia, stratifying patients by *SAMHD1* genotype or expression.

**Supplementary Figure 5.**
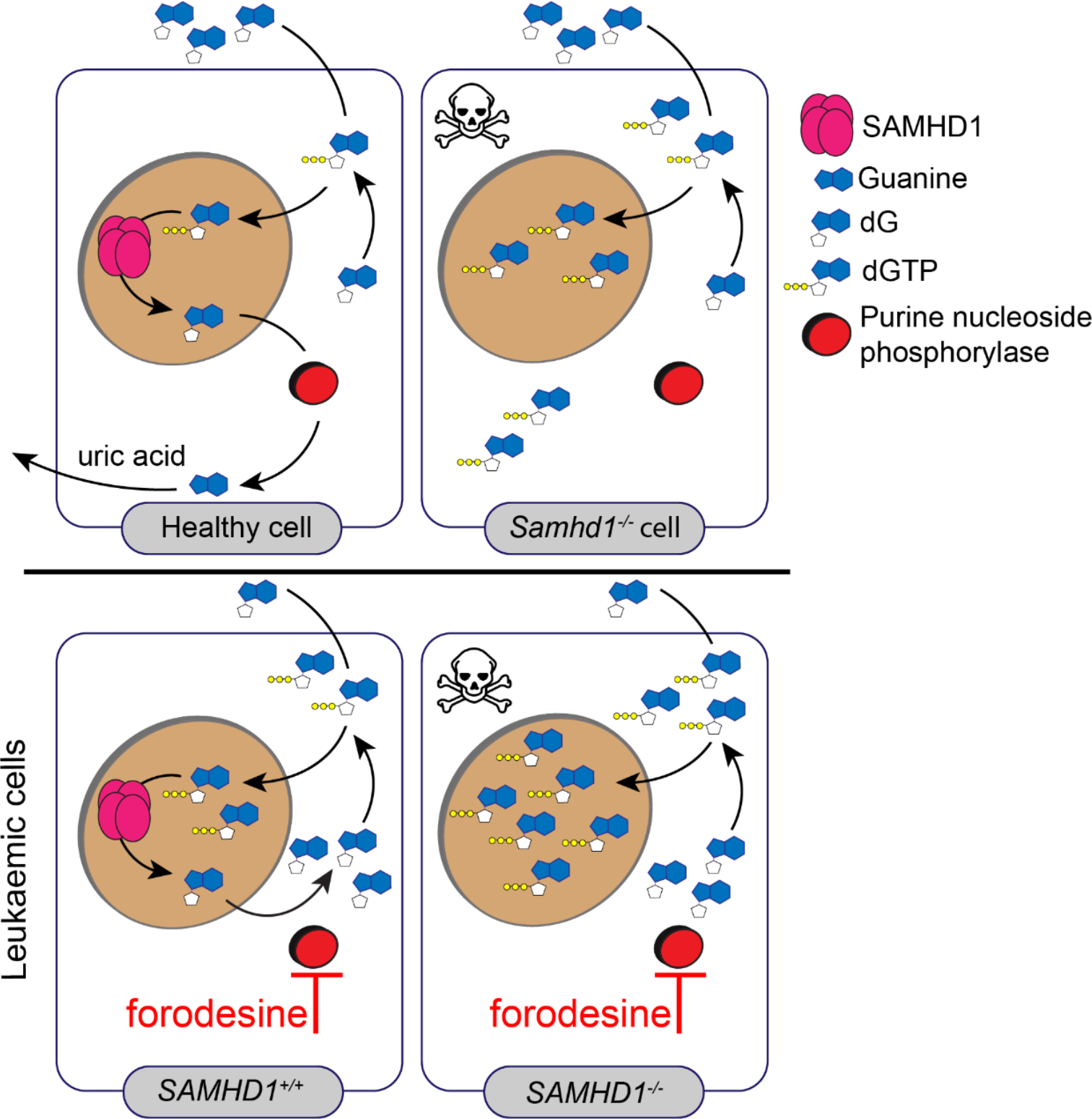
Graphical Abstract.

**Highlights:**

- SAMHD1-deficient cells die upon exposure to deoxyribonucleosides (dNs)
- Deoxyguanosine (dG) is the most toxic dN, inducing apoptosis in cells lacking SAMHD1
- *SAMHD1*-mutated leukaemic cells can be killed by dG and the PNP-inhibitor forodesine

**In Brief:** SAMHD1 degrades deoxyribonucleoside triphosphates (dNTPs), the building blocks of DNA. Davenne et al. found that SAMHD1 protects cells against dNTP imbalances. Exposure of SAMHD1-deficient cells to deoxyguanosine (dG) results in increased intracellular dGTP levels and subsequent apoptosis. This can be exploited to selectively kill cancer cells that acquired *SAMHD1* mutations.

## Introduction

Intracellular deoxyribonucleoside triphosphate (dNTP) concentrations are controlled by dNTP synthesis and degradation. dNTPs are supplied by two pathways known as de novo and salvage. In the de novo pathway, dNTPs are synthesised from intracellular precursors. The enzyme ribonucleotide reductase catalyses the rate limiting step and converts ribonucleoside diphosphates into deoxyribonucleoside diphosphates (Hofer et al., 2012). The salvage pathway involves uptake of deoxyribonucleosides (dNs) from the extracellular environment, followed by intracellular phosphorylation by cytosolic and mitochondrial kinases to form dNTPs (Eriksson et al., 2002; Inoue, 2017; Reichard, 1988).

One enzyme degrading intracellular dNTPs is the phosphohydrolase SAMHD1, initially identified as an interferon *γ*-inducible transcript in dendritic cells (Li et al., 2000). SAMHD1 cleaves all four dNTPs into the corresponding dNs and inorganic triphosphate (Goldstone et al., 2011; Powell et al., 2011). The catalytically active form of the protein is a homo-tetramer, formation of which is regulated allosterically by dNTPs and guanosine triphosphate (GTP) as well as by phosphorylation of threonine 592 (reviewed in: Ahn, 2016; Ballana and Este, 2015). SAMHD1 has been studied extensively in the context of human immunodeficiency virus (HIV) infection. By limiting the supply of dNTPs for the viral reverse transcriptase, SAMHD1 blocks HIV infection in certain cell types (Hrecka et al., 2011; Laguette et al., 2011; Lahouassa et al., 2012; Rehwinkel et al., 2013). *SAMHD1* mutations cause Aicardi-Goutières syndrome, a rare autoinflammatory disease characterised by chronic production of type I interferons, a family of cytokines typically upregulated only during acute virus infection (Crow and Manel, 2015; Rice et al., 2009). Furthermore, mutations in the *SAMHD1* gene have been found in several types of cancer, including colorectal cancer and leukaemias (Clifford et al., 2014; Johansson et al., 2018; Rentoft et al., 2016; Schuh et al., 2012). It is possible that inactivation of SAMHD1 provides transformed cells with a growth advantage simply due to elevated dNTP levels. Alternatively, the role of SAMHD1 in cancer may relate to its functions in DNA repair and DNA replication, which are independent of dNTP degradation (Clifford et al., 2014; Coquel et al., 2018; Daddacha et al., 2017).

Chronic lymphocytic leukaemia (CLL) is a very common form of adult leukaemia and affects the elderly (Swerdlow, 2008). Refractoriness to chemotherapy and relapse remain major causes of death of patients with CLL. Nucleotide metabolism is an attractive target for the treatment of CLL and other leukaemias. The small molecule forodesine (also known as Immucillin H or BCX-1777) was developed to inhibit purine nucleoside phosphorylase (PNP) (Kicska et al., 2001). PNP degrades deoxyguanosine (dG) into guanine, which is further catabolised into uric acid that is released by cells (Gabrio et al., 1956). dG has cytotoxic properties (Dahbo and Eriksson, 1985; Mann and Fox, 1986; Theiss et al., 1976) and genetic PNP deficiency causes immunodeficiency and results in loss of T cells and in some patients also affects B cell function (Markert, 1991). Upon forodesine treatment, dG accumulates intracellularly and is phosphorylated to deoxyguanosine triphosphate (dGTP). The resulting imbalance in dNTP pools is predicted to cause cell death and to eliminate leukaemic cells (Bantia et al., 2001). Furthermore, synergy between dG and forodesine in inducing cell death in vitro has been suggested (Bantia et al., 2003) and, in patients, forodesine treatment increases plasma dG levels (Balakrishnan et al., 2006; Balakrishnan et al., 2010). Forodesine showed promising results in vitro in killing CLL B cells; surprisingly, however, it had substantial activity only in a small subset of patients with B or T cell malignancies (Alonso et al., 2009; Balakrishnan et al., 2006; Balakrishnan et al., 2010; Dummer et al., 2014; Gandhi and Balakrishnan, 2007; Gandhi et al., 2005; Maruyama et al., 2018).

Here, we explored the role of SAMHD1 in dNTP metabolism. We report that SAMHD1 protected cells against imbalances in dNTP pools. In cells lacking SAMHD1, engagement of the salvage pathway resulted in programmed cell death. Exposure to dG was particularly potent at inducing intrinsic apoptosis in SAMHD1-deficient primary and transformed cells. We further show that forodesine and other PNP inhibitors acted synergistically with dG to induce death in cells lacking SAMHD1. Importantly, *SAMHD1*-mutated leukaemic cells from patients with CLL were selectively killed by forodesine and dG. This showed that SAMHD1 was limiting the potency of forodesine. It may therefore be possible to stratify patients with leukaemia for forodesine treatment by *SAMHD1* genotype or expression.

## Results

### SAMHD1 protects cells against dNTP overload

To investigate the role of SAMHD1 in dNTP metabolism, we added equimolar concentrations of dNs to wild-type (WT) or SAMHD1-deficient cells. Surprisingly, wide-spread cell death was apparent by bright-field microscopy in cells lacking SAMHD1, but not in control cells after over-night incubation with dNs (data not shown). To study this phenotype systematically, we analysed mouse embryonic fibroblasts (MEFs), mouse bone marrow-derived macrophages (BMDMs) and primary human fibroblasts. Cell viability was assessed using a luminescence-based assay for intracellular ATP levels (CellTiter-Glo). We observed reduced viability of dN-exposed *Samhd1^-/-^* MEFs and BMDMs, as well as of human fibroblasts from a patient with AGS homozygously carrying the Q149X nonsense mutation in *SAMHD1* (Figure 1A). Viability of WT mouse and control human cells, including fibroblasts from patients with AGS carrying other AGS-causing mutations in *IFIH1* or *ADAR1,* was largely unaltered after addition of dNs. To confirm that the absence of SAMHD1 renders cells susceptible to dN-induced cell death, we reconstituted BMDMs with a retrovirus expressing mouse SAMHD1. Indeed, expression of SAMHD1 in *Samhd1^-/-^* cells rescued viability after treatment with dNs (Figure 1B,C).

**Figure 1.**
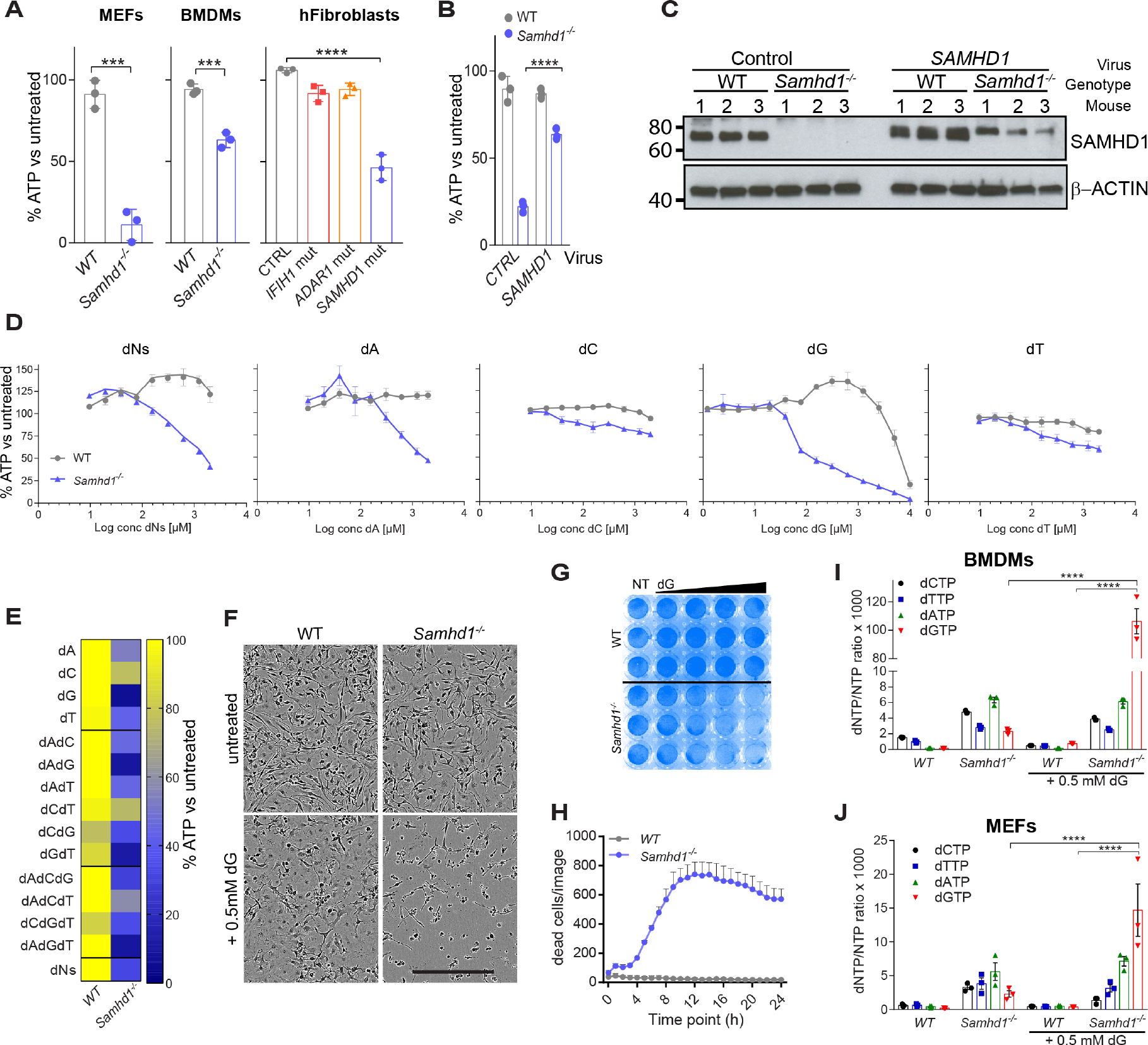
Deoxyribonucleosides (dNs) are toxic in SAMHD1-deficient cells. (**A**) MEFs, BMDMs and AGS patient-derived fibroblasts were treated with a mix of all four dNs. MEFs were cultured with 0.8 mM of each dN for 48 hours. BMDMs and fibroblasts were treated with 0.5 mM of each dN for 24 hours. Cell viability was determined by CellTiter-Glo assay. For each genotype, values from untreated control cells were set to 100%. Data from triplicate measurements are shown with mean ± SD. P-values determined with unpaired t-tests (MEFs and BMDMs) or one-way ANOVA (fibroblasts) are indicated. (**B,C**) SAMHD1 expression was reconstituted in *Samhd1^-/-^* BMDMs. Cells of the indicated genotype were infected with a retrovirus expressing SAMHD1 or empty control retrovirus. Cells were then treated with 0.5 mM of each dN for 24 hours. (**B**) Cell viability was tested as in (**A**). Values from triplicate measurements are shown with mean ± SD. P-values determined with two-way ANOVA are indicated. (**C**) SAMHD1 expression was tested by western blot in BMDMs from three mice per genotype. β-Actin served as a loading control. (**D**) BMDMs were treated with equimolar concentrations of all four dNs or with individual dNs at the indicated concentrations for 24 hours. Cell viability was tested as in (**A**). Data from biological triplicates are shown as mean ± SEM. (**E**) BMDMs were treated with individual dNs and combinations of dNs. Cells were cultured with 0.5 mM of the indicated dN(s) for 24 hours and viability was analysed as in (**A**). Data from biological triplicates were averaged and are represented as a heat map. (**F**) BMDMs were treated with 0.5 mM dG for 24 hours. Brightfield images are shown. The scale bar represents 300 µm. (**G**) BMDMs were treated with increasing doses of dG for 24 hours, fixed and stained with crystal violet. The wedge denotes 0.2, 0.4, 0.8, 1.6 mM dG; NT, not treated. (**H**) BMDMs were treated with 0.4 mM dG. Viability was monitored with the cell-impermeable dye Yoyo3 for 24 hours using an in-incubator imaging system (Incucyte). Yoyo3^+^ cells were enumerated. Mean values from triplicate measurements are shown ± SD. (**I,J**) BMDMs (**I**) and MEFs (**J**) were treated with 0.5 mM dG for 2 hours and intracellular dNTP levels were quantified relative to NTP levels. Data from three biological replicates are shown together with mean ± SEM. P-values determined with two-way ANOVA are indicated. Panels **A**-**C** and **F**-**H** are representative of at least three independent experiments. *** p<0.001; **** p<0.0001.

Next, we exposed BMDMs to increasing concentrations of dNs. We observed dose-dependent toxicity in *Samhd1^-/-^* cells, but not in WT cells, starting at ∼0.1 mM dNs (Figure 1D). To determine if this effect was due to a specific dN, we treated BMDMs with single dNs. Interestingly, the highest toxicity in *Samhd1^-/-^* cells was observed when dG was used (Figure 1D). Like dG treatment, dA also reduced viability specifically in SAMHD1-deficient cells, but at higher doses: a 50% reduction in intracellular ATP levels was observed with ∼0.1 mM dG and with ∼1 mM dA (Figure 1D). Of note, dG also caused toxicity in WT cells at high doses above 5 mM (Figure 1D). We also tested dN combinations using a fixed dose of 0.5 mM. dG was the most toxic dN in *Samhd1^-/-^* cells when used alone or in combination with dA and/or thymidine, whilst presence of dC reduced the effect of dG on cell viability (Figure 1E). We therefore focussed on dG in subsequent experiments at doses that do not reduce viability in WT cells. Bright-field images, crystal violet staining and live-cell imaging confirmed the toxicity of dG in *Samhd1^-/-^* cells (Figure 1F-H). In line with earlier work (Behrendt et al., 2013; Rehwinkel et al., 2013), measurement of intracellular dNTP concentrations showed that the levels of all four dNTPs were elevated in *Samhd1^-/-^* cells (Figure 1I,J). Importantly, dG treatment resulted in 46- and 6-fold increases in dGTP concentrations in *Samhd1^-/-^* BMDMs and MEFs, respectively, while dGTP levels stayed largely unchanged in WT cells (Figure 1I,J). Taken together, these data show that exposure to dG led to dGTP accumulation in cells lacking SAMHD1, subsequently resulting in cell death.

### dG treatment induces apoptosis in *Samhd1^-/-^* cells

We next determined the type of cell death triggered by dNs in *Samhd1^-/-^* cells. Annexin V and 7AAD staining showed an increased frequency of early apoptotic (AnnexinV^+^7AAD^-^) and dead (AnnexinV^+^7AAD^+^) cells in dN-treated *Samhd1^-/-^* BMDM cultures (Figure 2A). Using the Caspase-Glo assay to measure activity of apoptotic caspases, we found that addition of dG to *Samhd1^-/-^* BMDMs, but not to WT cells, activated caspase 3/7 (Figure 2B). Live cell imaging revealed that *Samhd1^-/-^* BMDMs treated with dG stained positive for Annexin V around 5 hours post-treatment and subsequently for propidium iodide (PI) (Figure 2C). These observations suggest that dG treatment induced apoptosis, followed by secondary necrosis rendering cells permeable to PI. Quantification of AnnexinV^+^PI^+^ cells by microscopy 24 hours after dG exposure affirmed increased levels of dead *Samhd1^-/-^* BMDMs (Figure 2D). Cleaved caspase 3 was detectable by western blot in *Samhd1^-/-^* but not WT BMDMs, supporting the notion that dG induced apoptosis (Figure 2E). Cycloheximide (CHX) served as a control in these experiments and induced caspase 3 cleavage in WT and *Samhd1^-/-^* cells. Cytochrome C is normally present in mitochondria and is released into the cytosol early during intrinsic apoptosis. To characterise the apoptosis pathway activated by dG-treatment, we measured cytochrome C levels in the cytosol and found that treatment of *Samhd1^-/-^* BMDMs with dG lead to a redistribution of cytochrome C into the cytosol (Figure 2F). To investigate whether a soluble factor was triggering apoptosis, we co-cultured WT and *Samhd1^-/-^* BMDMs and treated them with dG. Viability decreased with increasing proportions of *Samhd1^-/-^* cells present in the co-culture, suggesting that apoptosis was induced in a cell-autonomous fashion (Figure 2G). In sum, these results show that dG triggered intrinsic apoptosis in *Samhd1^-/-^* cells.

**Figure 2.**
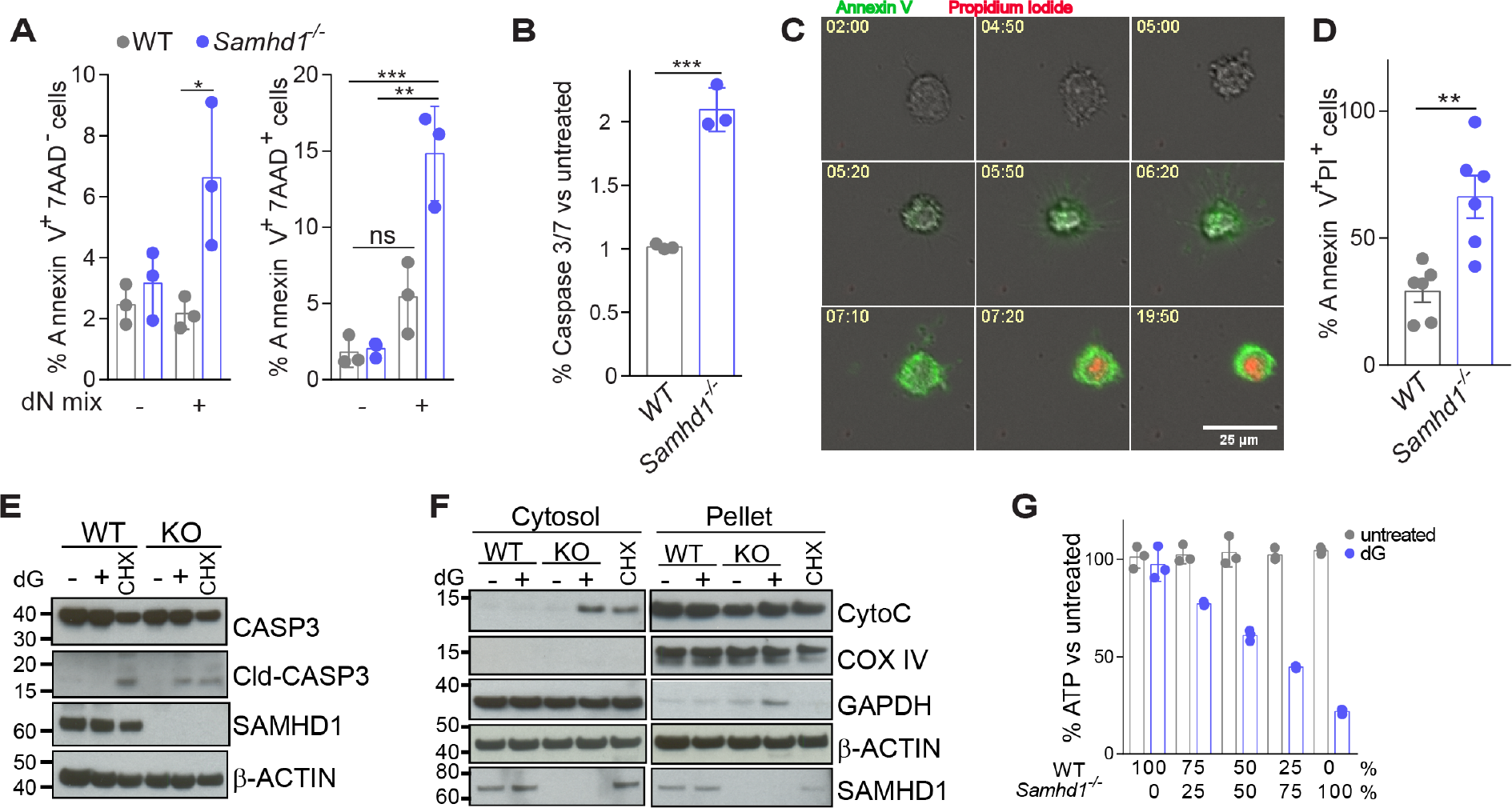
dG treatment kills *Samhd1^-/-^* cells by apoptosis. (**A**) BMDMs were treated with 0.5 mM of each dN for 24 hours and stained with Annexin V and 7AAD. AnnexinV^+^7AAD^-^ and AnnexinV^+^7AAD^+^ cells were quantified by flow cytometry. Data from triplicate measurements are shown with mean ± SD. P-values determined with two-way ANOVA are indicated. (**B**) Caspase activity was assessed in BMDMs 6 hours after treatment with 0.5 mM dG using the Caspase Glo 3/7 assay. For each genotype, values from untreated control cells were set to 1. Data from triplicate measurements are shown with mean ± SD. The p-value determined with an unpaired t-test is indicated. (**C,D**) Live cell imaging of *Samhd1^-/-^* BMDMs treated with 0.5 mM dG. Alexa-488-labelled Annexin V and propidium iodide (PI) were added to the culture medium to visualise early apoptotic cells and cells that lost membrane integrity, respectively. (**C**) Representative images. Numbers show the time after dG exposure (hh:mm). (**D**) Enumeration of AnnexinV^+^ PI^+^ cells after 24 hours of treatment with 0.5 mM dG. Six images per condition were analysed and means ± SEM are shown. The p-value determined with an unpaired t-test is indicated. (**E,F**) BMDMs were treated with 0.5 mM dG or 1 µg/ml cycloheximide (CHX, added to WT cells in [**F**]) for 8 hours. (**E**) Levels of the indicated proteins in total cell extracts were determined by western blot. (**F**) Cells were fractionated into cytosol and a pellet containing organelles. Levels of the indicated proteins were determined by western blot. *β*-Actin served as a loading control. (**G**) WT and *Samhd1^-/-^* BMDMs were co-cultured at the indicated ratios. Cell viability was determined as in Figure 1A 24 hours after treatment with 0.5 mM dG. Data from triplicate measurements are shown with mean ± SD. Panels **A-G** are representative of at least three independent experiments. ns p≥0.05; * p<0.05; ** p<0.01; *** p<0.001. See also Supplementary Figure 1.

### Nuclear DNA replication is not required for dG-induced apoptosis

Earlier work in yeast showed that severe dNTP pool imbalances can trigger stalled replication forks and checkpoint activation (Kumar et al., 2011; Poli et al., 2012). To study the role of DNA replication in dG-induced death of SAMHD1-deficient cells, we analysed cell cycle progression in BMDMs by BrdU and PI staining after dG treatment. Untreated WT and *Samhd1^-/-^* BMDM cultures contained ∼20% BrdU^+^ cells, indicative of cells in S-phase with ongoing DNA replication (Supplementary Figure 1A). After dG treatment, WT cells already in S-phase progressed through the cell cycle. At the same time, new cells did not enter S-phase, resulting in a much-reduced population of BrdU^+^ cells after 24 hours of dG exposure. In contrast, *Samhd1^-/-^* cells in S-phase did not progress. Instead, a population of cells displaying sub-G0/G1 PI staining, indicative of dead cells that lost their nucleic acid content, was detected in *Samhd1^-/-^* cultures, starting at 8 hours after dG treatment (Supplementary Figure 1A). Next, we performed a BrdU pulse-chase experiment, in which we labelled BMDMs with BrdU first and then treated with dG. Over time, WT cultures accumulated a distinct population of G0/G1-BrdU^+^ cells and contained fewer cells in S-phase (Supplementary Figure 1B). This confirmed that WT cells progressed through the cell cycle but did not enter S-phase. In *Samhd1^-/-^* cultures, cells with sub-G0/G1 PI staining were evident from 8 hours onwards. These included both BrdU^+^ and BrdU^-^ cells, suggesting that dG treatment killed both cycling and non-cycling *Samhd1^-/-^* cells (Supplementary Figure 1B). We therefore tested whether DNA replication was required for the toxicity of dG in *Samhd1^-/-^* cells. BMDMs were cultured in serum-free medium (R0) or were pre-treated with hydroxyurea (HU), both of which induced cell cycle arrest, evident from reduced numbers of cells in S-phase (Supplementary Figure 1C). *Samhd1^-/-^* cells arrested by both methods were susceptible to killing by dG (Supplementary Figure 1D,E). We also pre-treated BMDMs with aphidicolin (APD) that blocks nuclear but not mitochondrial DNA polymerases (Lentz et al., 2010; Zimmermann et al., 1980). As expected, APD-treated cells were arrested in early S-phase (Supplementary Figure 1F). Interestingly, APD-treated cells were susceptible to dG-induced toxicity (Supplementary Figure 1G). Finally, we assessed oxidative stress by measuring levels of the reactive oxygen species H_2_O_2_ and found that dG treatment of WT and SAMHD1-deficient BMDMs did not induce oxidative stress (Supplementary Figure 1H). As a control, we used menadione that induced equivalent H_2_O_2_ levels in cells irrespective of their genotype. Together, these data suggest that dG-induced apoptosis occurred independently of nuclear DNA replication and oxidative stress, and that dGTP overload was toxic in both cycling and non-cycling *Samhd1^-/-^* cells.

### dG treatment kills SAMHD1-deficient cancer cells

*SAMHD1* mutations are present in several types of cancer and in many cases result in reduced mRNA and protein levels (Clifford et al., 2014; Johansson et al., 2018; Rentoft et al., 2016). We therefore wished to explore our finding of dN-induced cell death in the context of malignant disease. Initially, we tested cancer cell lines. Vpx is a HIV-2 accessory protein that targets SAMHD1 for proteasomal degradation (Hrecka et al., 2011; Laguette et al., 2011). We used virus-like particles (VLPs) containing Vpx to deplete SAMHD1 in the cervical cancer cell line HeLa, in the monocytic cell line THP1 and in the breast cancer cell line MDA-MB231, all of which express SAMHD1. Cells treated with VLPvpx, but not with control VLPs lacking Vpx (VLP_ctrl_), showed reduced viability upon addition of dNs or dG (Figure 3A-D). SAMHD1 staining and analysis by flow cytometry confirmed the efficiency of SAMHD1 depletion using VLPvpx (Supplementary Figure 2A,B). In addition, we generated a *Samhd1^-/-^* B16F10 mouse melanoma cell line using CRISPR/Cas9 (strategy and validation shown in Supplementary Figure 2C,D). *Samhd1^-/-^* B16F10 cells showed increased frequencies of early apoptotic (AnnexinV^+^7AAD^-^) and dead (AnnexinV^+^7AAD^+^) cells upon dG treatment, accompanied by reduced confluency (Figure 3E-G and Supplementary Figure 2E). We made similar observations in the murine colorectal cancer cell line CT26 upon SAMHD1 knock-out (data not shown). We also included Jurkat cells in our analysis, a human T-cell line that, in contrast to the other cell lines utilized, does not express SAMHD1 (Baldauf et al., 2012). Jurkat cells were exquisitely sensitive to dG treatment at doses approximately 10 times lower than those used in most other experiments (Figure 3H). We confirmed the killing of Jurkat cells by testing their clonogenic potential, which was greatly reduced upon dG treatment (Figure 3I). Reconstitution with a lentivirus expressing human SAMHD1 partially rescued viability upon dG treatment (Figure 3J,K). Interestingly, SAMHD1 K11A, which does not localise to the cell nucleus (Schaller et al., 2014), executed an even more pronounced rescue as compared to WT SAMHD1. In sum, SAMHD1 protected cancer cell lines against dN-triggered toxicity.

**Figure 3.**
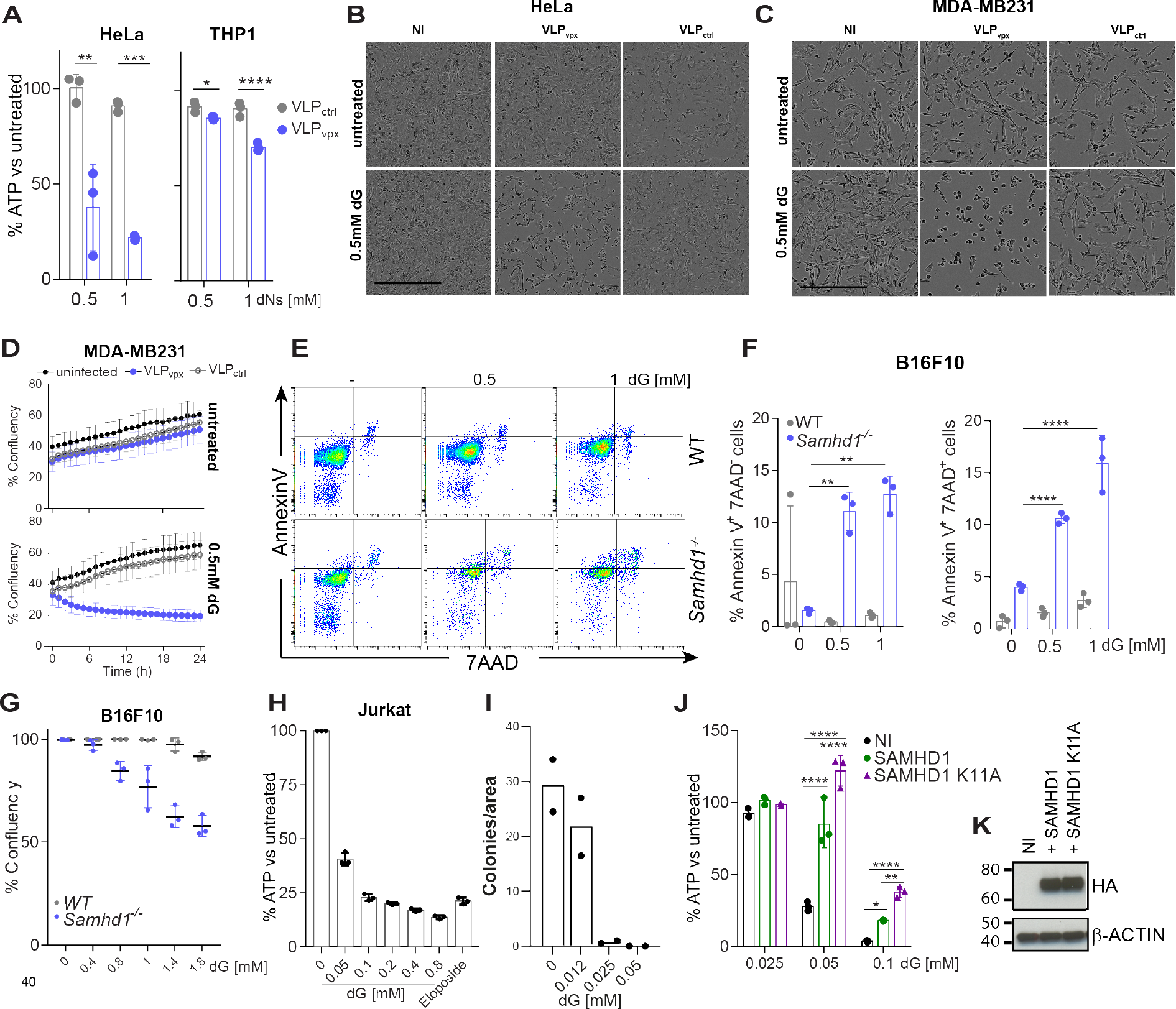
dG induces death of cancer cell lines. (**A**) HeLa and THP1 cells were infected with VLPs containing Vpx (VLP_vpx_) or not (VLP_ctrl_). After 24 hours, cells were treated with 0.5 mM of each dN for an additional 24 hours. Cell viability was assessed as in Figure 1A. (**B,C**) HeLa and MDA-MB231 cells were left un-infected (NI) or were infected with VLPs containing Vpx (VLP_vpx_) or not (VLP_ctrl_). After 24 hours, cells were treated with 0.5 mM dG and brightfield images were acquired after an additional 10-12 hours. Scale bar represents 300 µm. (**D**) MDA-MB231 were treated as in (**C**) and confluency was monitored after dG addition using a live-cell imaging system in the incubator (Incucyte). The mean of 9 measurements ± SD is shown. (**E-G**) Wild-type and *Samhd1^-/-^* B16F10 cells were treated with dG as indicated for 20 hours. (**E,F**) Cells were then stained with annexin V and 7AAD and analysed by flow cytometry. Representative FACS plots are shown in panel (**E**) and Annexin V^+^ 7AAD^-^ and Annexin V^+^ 7AAD^+^ cells were quantified in panel (**F**). (**G**) Confluency was determined as in panel (**D**). (**H**) Jurkat cells were treated for 20 hours with dG as indicated or with 25 µM etoposide. Cell viability was determined as in Figure 1A. (**I**) Jurkat cells were treated with dG as indicated for 20 hours and were then seeded in semi-solid medium containing dG. After 13 days, cell colonies were counted and the number colonies per field of view is shown. (**J,K**) Jurkat cells were reconstituted with HA-tagged wild-type or K11A mutant SAMHD1 using a lentivector. Un-infected cells (NI) served as control. (**J**) Cells were then treated with dG for 48 hours. Cell viability was determined as in Figure 1A. (**K**) SAMHD1 levels in total cell extracts were determined by western blot. β-Actin served as a loading control. Panels **A**, **D-H** and **J-K** are representative of three independent experiments and panels **B** and **C** of two experiments. In panels **A**, **F-H** and **J** dots represent technical triplicates and means ± SD are shown. In panel **I**, data from two independent experiments were pooled and dots represent the mean of technical duplicates per experiment. P-values determined with two-way ANOVA are indicated. * p<0.05; ** p<0.01; *** p<0.001; **** p<0.0001. See also Supplementary Figure 2.

### SAMHD1 protects against combined forodesine and dG treatment

Forodesine is an inhibitor of PNP, which converts dG into guanine and *α*-D-ribose 1-phosphate, and has been shown to induce apoptosis in leukaemic cells, possibly by increasing the intracellular and plasma concentrations of dG and consequently intracellular dGTP (Balakrishnan et al., 2010; Bantia et al., 2010; Kicska et al., 2001; Posmantur et al., 1997). Our system – in which SAMHD1-deficient cells fed with dG died by apoptosis due to dGTP overload – resembled this situation. We therefore hypothesised that forodesine and dG might work synergistically in SAMHD1-deficient cells. Indeed, low doses of dG or forodesine alone did not compromise viability of WT or *Samhd1^-/-^* BMDMs while the combination of both was toxic in *Samhd1^-/-^* cells (Figure 4A,B). We confirmed this observation using 10 µM dG and 1 µM forodesine at (i) different time points, (ii) with crystal violet staining and (iii) biochemically by cleavage of PARP and CASPASE3 (Figure 4C-F). Jurkat cells treated with the same doses of forodesine and dG showed increased proportions of AnnexinV^+^7AAD^-^ cells, and this was prevented when SAMHD1 or SAMHD1 K11A were expressed (Figure 4G-I). Vice versa, SAMHD1-sufficient Hela cells were sensitised to combined forodesine and dG treatment by Vpx-mediated depletion of SAMHD1 (Figure 4J).

**Figure 4.**
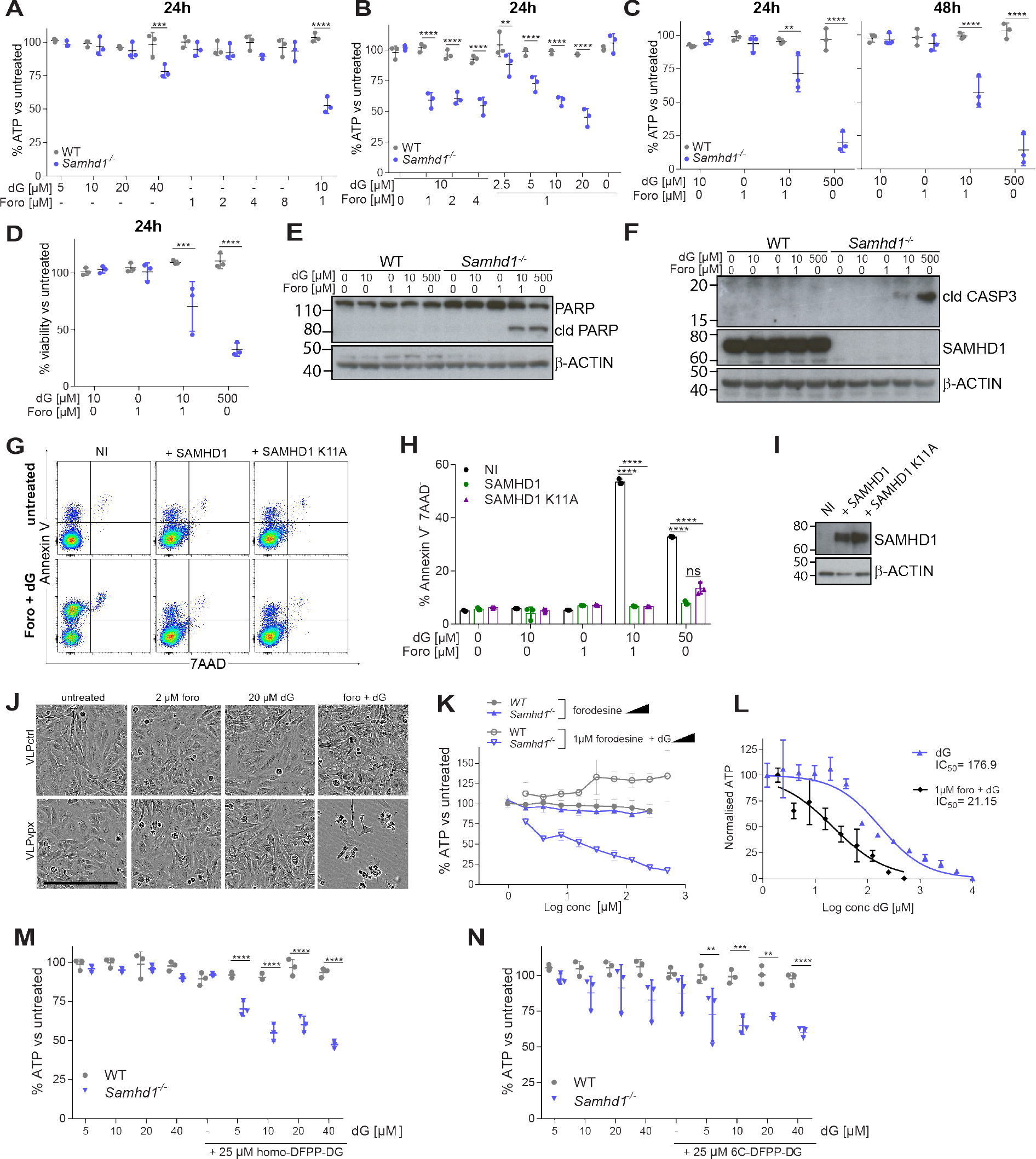
PNP inhibitors and dG synergistically induce cell death in cells lacking SAMHD1. (**A-C**) BMDMs were treated with the indicated doses of dG and Forodesine. Viability was tested as in Figure 1A after 24 or 48 hours. (**D**) BMDMs treated for 24 hours with dG and Forodesine were fixed and stained with crystal violet. After washing, cell-associated dye was solubilised and quantified by absorbance at 570 nm. For each genotype, values from untreated control cells were set to 100%. (**E-F**) BMDMs were treated for 8 hours with dG and Forodesine. Levels of the indicated proteins in total cell extracts were determined by western blot. β-Actin served as a loading control. cld, cleaved. (**G-I**) Jurkat cells were reconstituted with SAMHD1 as described in Figure 3J,K. Un-infected cells (NI) served as control. (**G,H**) Cells were treated for 18 hours with 10 µM dG and 1 µM forodesine. Cells were then stained with annexin V and 7AAD and analysed by flow cytometry. Representative FACS plots are shown in panel (**G**) and Annexin V^+^ 7AAD^-^ cells were quantified in panel (**H**). (**I**) SAMDH1 levels in total cell extracts were determined by western blot. *β*-Actin served as a loading control. (**J**) HeLa cells were infected with VLPs containing Vpx (VLP_vpx_) or not (VLP_ctrl_). After 6 hours, cells were treated with 20 µM dG and 2 µM forodesine and brightfield images were acquired after an additional 48 hours. Scale bar represents 300 µm. (**K**) BMDMs were treated with the indicated doses of dG and forodesine. Viability was tested as in Figure 1A after 24 hours. Means from three biological replicates are shown ± SEM. (**L**) *Samhd1^-/-^* BMDMs were treated with the indicated doses of dG in the presence or absence of 1 µM forodesine. Cell viability was determined by CellTiter-Glo assay after 24 hours. Data were normalised by setting the values for the lowest and highest dG concentrations to 100 and 0, respectively. Means from three biological replicates are shown ± SEM. IC50 values were calculated from the non-linear regression curves shown on the graph. (**M**,**N**) BMDMs were treated with the indicated doses of dG and homo-DFPP-DG (**M**) or 6C-DFPP-DG (**N**). Viability was tested as in Figure 1A after 24 hours. Data are representative of three independent experiments. In panels **A-D, M, N** and **H** dots represent BMDMs from individual mice and technical replicates, respectively. Mean ± SD is shown. P-values determined with two-way ANOVA are indicated. ** p<0.01; *** p<0.001; **** p<0.0001

To further study the synergy between forodesine and dG in cells lacking SAMHD1, we titrated forodesine and dG. Forodesine alone had no effect on viability of *Samhd1^-/-^* BMDMs, including at high doses (Figure 4K, closed symbols). In contrast, dG triggered dose-dependent toxicity in the presence of 1 µM forodesine in *Samhd1^-/-^* cells but not in WT cells (Figure 4K, open symbols). Comparing the dose-response to dG in the presence or absence of forodesine revealed that forodesine sensitised SAMHD1-deficient BMDMs to dG by approximately 10-fold (Figure 4L).

Finally, we tested whether other PNP inhibitors might induce death of SAMHD1-deficient cells in the presence of dG. Indeed, BMDMs showed reduced viability upon exposure to either homo-DFPP-DG or 6C-DFPP-DG (Glavas-Obrovac et al., 2010; Hikishima et al., 2007, 2010) together with low doses of dG (Figure 4M,N). Taken together, these data showed that SAMHD1 protected cells against death that was synergistically induced by PNP inhibitors and dG. Thus, our observations revealed a key role of SAMHD1 in the mechanism underlying the toxicity of compounds such as forodesine.

### CLL B cells with *SAMHD1* mutations are highly sensitive to a combination treatment of forodesine and dG

*SAMHD1* is mutated in 11% of patients with refractory CLL (Clifford et al., 2014). Since SAMHD1 protected cells against treatment with forodesine and dG (Figure 4), we hypothesised that CLL B cells with *SAMHD1* mutations would be particularly susceptible to this combination treatment. To test this, we compared the effect of forodesine and dG treatment on peripheral blood mononuclear cells (PBMCs) from patients with CLL with or without acquired mutations in *SAMHD1*.

Details of the patients’ cells genetic status and *SAMHD1* mutations are shown in Supplementary Figure 3A,B. PBMCs from patients with CLL were treated with 2 µM forodesine, 20 µM dG, or both. When used alone, neither forodesine nor dG significantly reduced cell viability assessed by intracellular ATP content (Figure 5A). The combination of both compounds had little effect on viability of PBMCs from patients without *SAMHD1* mutations. However, significantly reduced PBMC viability was observed in the *SAMHD1*-mutated group (Figure 5A). These data were confirmed by flow cytometry: the population of live cells was selectively reduced after forodesine and dG treatment of PBMCs from *SAMHD1* mutated patients (data not shown).

**Figure 5.**
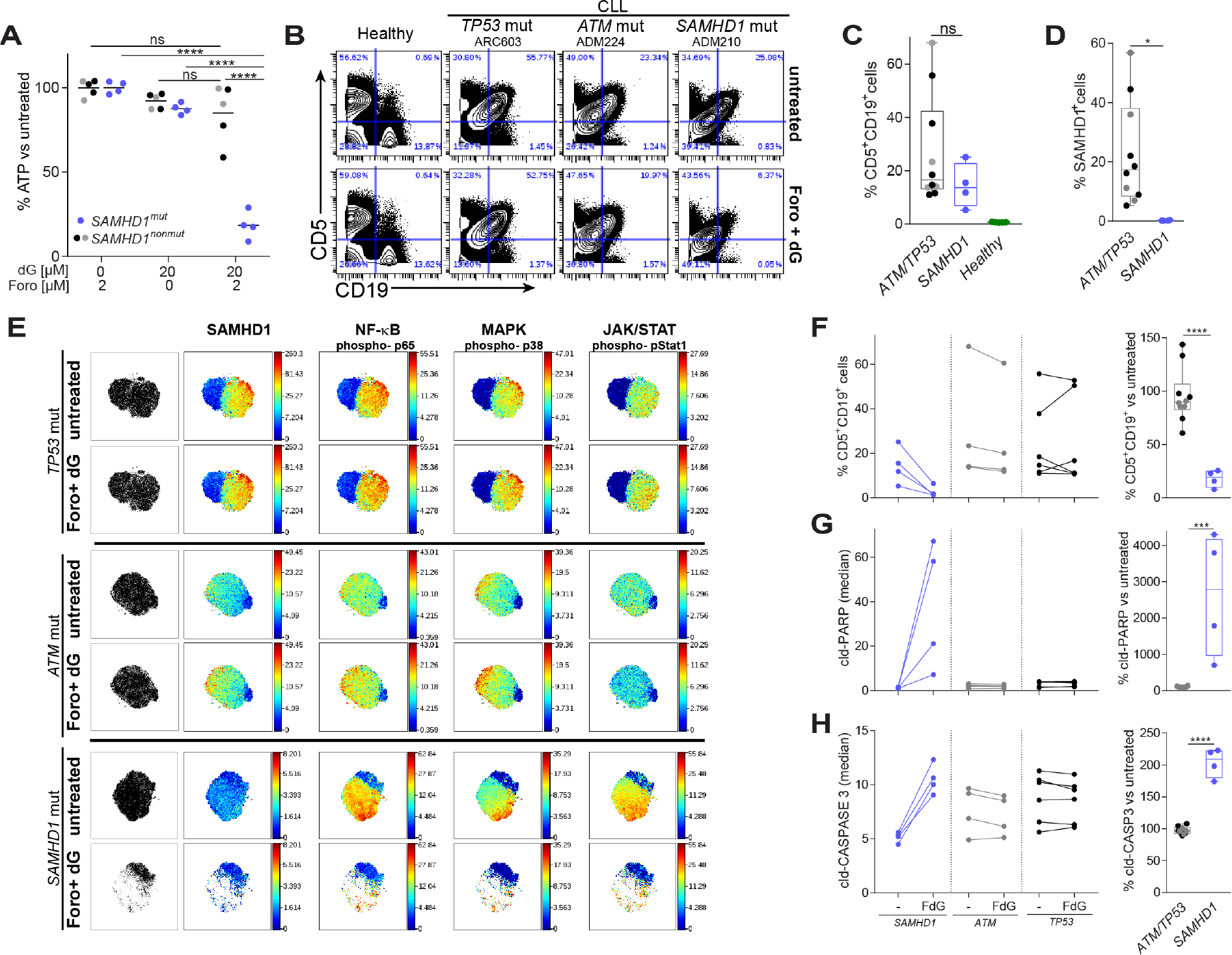
Elimination of *SAMHD1* mutated leukemic cells by Forodesine and dG treatment. (A) PBMCs from patients with CLL were treated for 24 hours with dG and Forodesine as indicated. Viability was tested as in Figure 1A. (**B-H**) PBMCs from healthy control subjects and patients with CLL were treated or not for 24 hours with 20 µM dG and 2 µM Forodesine (Foro + dG). Cells were then analysed using CyTOF. (B) Live cells were gated (see Supplementary Figure 3C). The CD5 and CD19 staining is shown for selected samples (see Supplementary Figure 3D for all samples). (C) Percentages of untreated, live CD5^+^CD19^+^ cells are shown. (D) SAMHD1 expression was analysed in untreated, live CD5^+^CD19^+^ cells and the percentage of SAMHD1^+^ cells is shown (see Supplementary Figure 3C for gating). (E) Live CD5^+^CD19^+^ cells from each sample were analysed separately by viSNE using 22 lineage markers (Cytobank; settings: 1000 iterations, 30 perplexity and 0.5 theta). Representative tSNE plots are shown (see Supplementary Figure 4 for all samples) and were coloured by expression or phosphorylation of the indicated markers. (F) Left, percentages of CD5^+^CD19^+^ cells amongst all live cells are shown in untreated and treated PBMC samples. FdG, treatment with 2 µM forodesine and 20 µM dG. Right, the frequency of live CD5^+^CD19^+^ cells was set to 100 in untreated samples and their percentage after forodesine and dG treatment is shown. (**G,H**) The staining for cleaved PARP (**G**) and cleaved CASPASE3 (**H**) in live CD5^+^CD19^+^ cells was analysed. Left, median values are shown in untreated and treated cells. Right, median values from untreated cells were set to 100 separately for each sample. In panels **A**, **C**, **D** and **F-H** dots represent cells from different patients and the colour indicates the mutation status (grey: *ATM*, black: *TP53*, blue: *SAMHD1*). Horizontal bars represent means and in panels **C** and **D** box and whiskers show SD and maximum/minimum values, respectively. P-values determined with two-way ANOVA (**A**) or unpaired t-test (**C**,**D**,**F-H**) are indicated. ns p≥0.05; * p<0.05; ** p<0.01; *** p<0.001; **** p<0.0001. See also Supplementary Figures 3-5.

To investigate which types of cells were affected by exposure to forodesine and dG, we used cytometry by time of flight (CyTOF) analysis. PBMCs from healthy control subjects and patients with CLL were treated or not with both compounds. After treatment, cells were stained with a panel of antibodies recognising cell surface markers to identify cell types. SAMHD1 expression, phosphorylation of NFκB-p65, p38 and STAT1, as well as cleavage of PARP and CASPASE3 were also monitored by intracellular staining. CLL B cells are marked by co-expression of CD5 and CD19 (Swerdlow, 2008). As expected, CD5^+^CD19^+^ cells (CLL B cells) were largely absent from control PBMCs and could be detected at varying frequencies in samples from patients with CLL, irrespective of *SAMHD1* genotype (Figure 5B,C and Supplementary Figure 3C,D). We also analysed SAMHD1 expression in CLL B cells. Variable expression of SAMHD1 was observed in the *SAMHD1* non-mutated group, whilst CLL B cells from the *SAMHD1* mutated group had no detectable levels of SAMHD1 (Figure 5D and Supplementary Figure 3C). This shows that the *SAMHD1* mutations studied here resulted in a loss of SAMHD1 protein, in line with our earlier observations (Clifford et al., 2014).

Next, we analysed our CyTOF data using viSNE (Amir el et al., 2013; Kimball et al., 2018). This analysis tool uses the t-distributed stochastic neighbour embedding (tSNE) algorithm and displays high-dimensional data on a two-dimensional map. Each dot on a viSNE plot corresponds to a cell. Colour can be used to show expression of a chosen parameter. We generated viSNE plots after gating on CLL B cells (Figure 5E and Supplementary Figure 4). tSNE maps showed a marked reduction of CLL B cells upon forodesine and dG treatment in cells from *SAMHD1*-mutated patients but not in the *SAMHD1* wild-type groups (Figure 5E and Supplementary Figure 4). Gating on CLL B cells also confirmed that forodesine and dG treatment resulted in their selective loss in the *SAMHD1*-mutated group while there was no effect on the same cell population in the *SAMHD1* non-mutated group (Figure 5F). Significantly increased levels of cleaved PARP and cleaved CASPASE3 were observed only in *SAMHD1-* mutated CLL B cells post-treatment, consistent with the induction of apoptosis (Figure 5G,H).

Interestingly, forodesine and dG ablated a subpopulation of *SAMHD1* mutated CLL B cells, which appeared to be characterised by high levels of NFκB-p65, p38 and STAT1 phosphorylation (Figure 5E and Supplementary Figure 4). Using phosphorylated (p-) p38, a MAP kinase, we defined “active” and “inactive” cells and analysed the expression of selected markers in these subpopulations (Supplementary Figure 5A). This analysis confirmed that p-p38 positive cells also displayed higher levels of p-p65 and p-STAT1 and revealed higher levels of CD27 expression in these “active” cells, which may indicate engagement of the B cell receptor (Lafarge et al., 2015). Interestingly, staining for cleaved PARP and cleaved CASPASE3 was enhanced more strongly in “inactive” cells upon treatment (Supplementary Figure 5B,C). This suggests that these “inactive” CLL B cells were also affected by the treatment and induced apoptosis with delayed kinetics compared to “active” cells. Collectively, these data show that forodesine and dG were highly efficient at killing malignant CLL B cells with *SAMHD1* mutations that cause a defect in SAMHD1 expression, while cells with intact SAMHD1 expression were spared.

## Discussion

Our data reveal an unexpected role of SAMHD1 in safeguarding cells against cell death resulting from imbalances in dNTP pools. In the absence of SAMHD1, dNTP imbalances induced by exposure of cells to dNs triggered apoptosis. This phenotype was observed in a wide range of human and mouse cells, including primary and transformed cells. We further show synergy between dG exposure and forodesine, which blocks dG degradation by PNP, in cells lacking SAMHD1. Importantly, the combination of dG and forodesine selectively killed SAMHD1-deficient CLL B cells, while other normal cells or SAMHD1-sufficient CLL B cells remained unaffected.

Since its identification as a restriction factor for HIV (Hrecka et al., 2011; Laguette et al., 2011), SAMHD1 has been studied extensively in the context of lentiviral infections. Interestingly, SAMHD1 is highly conserved from marine invertebrates to man (Rice et al., 2009), whereas lentiviruses evolved much more recently. This suggests that restriction of lentiviral infection is an exaptation of SAMHD1’s biochemical activity to degrade dNTPs, which perhaps has a more ancestral function in cellular dNTP metabolism. We propose that this evolutionarily conserved function of SAMHD1 is to correct imbalances in dNTP pools, thereby safeguarding against cell death. Indeed, we report that intracellular dNTP concentrations were only marginally altered in WT cells exposed to extracellular dNs, while SAMHD1-deficient cells accumulated large dNTP pools. Concomitantly, cells lacking SAMHD1 succumbed to apoptotic cell death. Interestingly, dG was the most toxic dN. This may be related to the observation that baseline dGTP concentrations are lower than those of the three other dNTPs, resulting in particularly pronounced dNTP imbalances upon dG feeding. In addition, dGTP allosterically regulates ribonucleotide reductase, preventing dCTP production through the de novo pathway (Moore and Hurlbert, 1966). Consistent with this idea is our observation that the toxicity of dG was reduced when added together with dC, providing dCTP via the salvage pathway. In addition, dC may indirectly increase dTTP concentrations via a pathway involving dCMP deaminase and thymidylate synthase (Theiss et al., 1976), thereby balancing dNTP levels.

The precise molecular mechanism by which dNTP imbalances in SAMHD1-deficient cells result in apoptosis remains to be fully elucidated. We observed release of cytochrome C from mitochondria into the cytosol, indicative of cell-intrinsic apoptosis. This notion was supported by the observation that cell death in mixed cultures containing WT and *Samhd1^-/-^* cells was proportional to the fraction of knock-out cells.

We further found that dN-triggered cell death did not require ongoing nuclear DNA synthesis. It is therefore possible that dNTP imbalances disrupt replication or repair of mitochondrial DNA, resulting in mitochondrial stress and subsequent apoptosis (Arpaia et al., 2000; Franzolin et al., 2015). Alternatively, dGTP may be involved more directly in the activation of apoptosis as has been reported for dATP (Li et al., 1997; Reubold et al., 2009).

Clinical trials showed that forodesine has beneficial effects in some but not all patients with B or T cell malignancies (Alonso et al., 2009; Balakrishnan et al., 2006; Balakrishnan et al., 2013; Balakrishnan et al., 2010; Dummer et al., 2014; Gandhi and Balakrishnan, 2007; Gandhi et al., 2005; Maruyama et al., 2018; Ogura et al., 2012), an observation that thus far has lacked an explanation. Our data suggest that forodesine-sensitive leukaemias harbour mutations that ablate SAMHD1 expression or inactivate its enzymatic activity. It may therefore be possible to stratify patients by *SAMHD1* genotype or expression levels. Retrospective analysis of previous clinical trials with forodesine could lend support for this idea. However, restricted sample availability, limited consent to obtain genetic information and small trial sizes have precluded this approach. Instead, future clinical trials should be conducted to determine whether the efficacy of forodesine can be predicted by the presence or absence of SAMHD1 in transformed cells. Survival of PBMCs ex vivo upon forodesine and dG treatment (Figure 5A) and commercially available *α*-SAMHD1 antibodies would be suitable for rapid clinical assays to identify patients with SAMHD1 deficiency. Previous studies reported that dG needs to be added in combination with forodesine to induce toxicity in vitro (Alonso et al., 2009; Balakrishnan et al., 2006; Bantia et al., 2001; Gandhi and Balakrishnan, 2007; Gandhi et al., 2005). In patients, forodesine treatment alone has been shown to increase plasma dG levels (Balakrishnan et al., 2006; Balakrishnan et al., 2010). However, it may also be interesting to explore supplementing forodesine with dG in patients with leukaemia with acquired *SAMHD1*-mutations. It is noteworthy that SAMHD1 is broadly expressed in most normal human tissues (Schmidt et al., 2015; Uhlen et al., 2015). As such, forodesine and dG are unlikely to have negative effects on healthy cells.

Our CyTOF analysis revealed that CLL B cells identified by CD19 and CD5 staining contained two sub-populations of cells, distinguishable by expression of CD27 and phosphorylation of p65, p38 and STAT1. Activation of NFκB, MAP kinase and STAT signalling pathways has been reported in CLL (Frank et al., 1997; Herishanu et al., 2011; Ringshausen et al., 2004; Shukla et al., 2018) and CD27 is upregulated in response to B cell receptor engagement (Lafarge et al., 2015). We therefore labelled these cell populations “active” and “inactive”. Interestingly, we find that forodesine and dG not only killed the “active” population of CLL B cells, but also induced markers of apoptosis in the “inactive” cells. As such, forodesine may have an advantage over other CLL drugs that inhibit B cell receptor signalling and thus target only “active” cells. In an independent line of investigations, SAMHD1 was found to not only degrade naturally occurring dNTPs but also some nucleotide analogues, including cytarabine (ara-C) and decitabine (DAC) that are used for treatment of acute myeloid leukaemia (AML) (Herold et al., 2017a; Herold et al., 2017b; Hollenbaugh et al., 2017; Oellerich et al., 2019; Schneider et al., 2017). The response of patients with AML to ara-C or DAC inversely correlates with SAMHD1 expression levels (Herold et al., 2017a; Oellerich et al., 2019; Schneider et al., 2017). These observations are an interesting parallel to our work and highlight SAMHD1 as a target for cancer therapy.

In summary, we uncovered an important role of SAMHD1 in protecting cells against dNTP imbalance that otherwise triggers apoptotic cell death. These findings allowed us to selectively ablate *SAMHD1*-mutated transformed cells using a combination treatment involving forodesine and dG. In future, forodesine may be developed into a precision medicine for a subset of patients with leukaemia with acquired *SAMHD1* mutations.

## Methods

### Plasmids

To generate pMSCVpuro-mSAMHD1, mouse *Samhd1* isoform 1 was PCR amplified. A kozak sequence and N-terminal 3xFLAG-tag were introduced by PCR and the PCR product was cloned into pMSCVpuro using the EcoRI site. To generate SAMHD1-deficient mouse cells, pX458-Ruby-sgRNA-1 and pX458-sgRNA-2 were cloned using pX458-Ruby and pX458, respectively, as described before (Hertzog et al., 2018).

### Mice

*Samhd1^-/-^* mice (C57BL/6 background) were described previously (Rehwinkel et al., 2013). 8-12 week old, male and female C57BL/6 WT and *Samhd1^-/-^* mice were used to obtain bone marrow for BMDM cultures.

### Cell culture

MEFs were made by standard protocols from either WT or *Samhd1^-/-^* mice. Bone marrow cells were isolated by standard protocols and, to obtain BMDMs, were grown in petri dishes for 7 days in R10 medium [Roswell Park Memorial Institute 1640 (RPMI) medium, 10% heat-inactivated foetal calf serum (FCS), 100 U/ml penicillin and 100 µg/ml streptomycin (P/S), 2 mM L-Glutamine] supplemented with 20% L929 conditioned medium and used on day 7. Human fibroblasts from AGS patients were provided by Y. Crow and G.I. Rice. MEFs were cultured in D10 medium [Dulbecco’s modified Eagle medium (DMEM) containing 10% heat-inactivated FCS, P/S, 2mM L-Glutamine and 20 mM HEPES buffer]. Human fibroblasts, HeLa, B16F10 and MDA MB231 cells were cultured in D10 without P/S. HeLa cells were a gift from M. Way, MDA MB231 cells were from A. Banham and B16F10 cells were provided by V. Cerundolo. Jurkat cells were a gift from S. Davis and originate from the American Type Tissue Collection and were cultured in R10 without P/S. All cells were cultured under 5% CO_2_. Human fibroblasts and MEFs were cultured under low oxygen (1.2% O_2_).

### Samples from patients with CLL

PBMCs from 19 patients with CLL recruited into the ADMIRE (n=12) and ARCTIC (n=7) studies (Howard et al., 2017; Munir et al., 2017) were retrieved from the Liverpool Bio-Innovation Hub Biobank. Genetic characterisation of the tumour cells for these patients was previously published (Clifford et al., 2014). PBMCs were thawed in R10 with P/S and 50 U/ml of benzonase, washed twice before being counted and plated. For CellTiter-Glo assay, 50,000 cells were plated in U-bottom 96-well plates. CyTOF experiments were performed using 3,000,000 cells in 12-well non-coated tissue culture plates.

### dNTP measurements

Cells from 4 plates (90 × 15 mm) of BMDMs or 3 plates (150 mm × 20 mm) of MEFs were pooled for each sample. Measurements were done on cells from different mice. Cells were treated with deoxynucleosides for a specific time and washed twice with ice-cold NaCl (9 g/L) on ice. Cells were then scraped in 550 µL of ice-cold trichloroacetic acid (15% w/v), MgCl2 (30mM) solution, collected into an Eppendorf tube, frozen on dry ice and stored at −80 °C. Cells were thawed on ice and processed as described in (Kong et al., 2018). Briefly, the cell suspension was pulse-vortexed (Intellimixer) at 99 rpm for 10 min at 4°C and centrifuged at 20,000 × g for 1 min at 4°C. The resulting supernatant was then neutralized twice with Freon-Trioctylamine mix (78% v/v - 22%, v/v respectively) by vortexing for 30 sec and centrifugation at 20,000 × g for 1 min. 475 µL of the aqueous phase was pH-adjusted with 1 M ammonium carbonate (pH 8.9), loaded on boronate columns (Affi-Gel Boronate Gel; Bio-Rad), and eluted with a 50 mM ammonium carbonate (pH 8.9) and 15 mM MgCl2 mixture to separate dNTPs from NTPs. The eluates containing dNTPs were adjusted to pH 4.5 and loaded onto an Oasis weak anion exchange (WAX) SPE cartridge. Interfering compounds were eluted off the cartridges in two steps with 1 mL ammonium acetate buffer (pH-adjusted to 4.5 with acetic acid) and 1 mL 0.5% ammonia aqueous solution in methanol (v/v), and the analytes were eluted from the cartridge with 2 mL methanol/water/ammonia solution (80/15/5, v/v/v) into a glass tube and then evaporated to dryness using a centrifugal evaporator at a temperature below 37°C. The residue was reconstituted in 1250 µL ammonium bicarbonate buffer, pH-adjusted to 3.4 and used for the HPLC analysis as described in (Jia et al., 2015). Briefly, nucleotides were isocratically eluted using 0.36 M ammonium phosphate buffer (pH 3.4, 2.5 % v/v acetonitrile) as mobile phase. dNTPs were normalized to total NTP pool of the cells.

### Viability assays

CellTiter-Glo (Promega), a luminescence assay that measures ATP levels, was used according to manufacturer instructions to assess viability. To assess cell viability with crystal violet, cells were stained with 0.5% crystal violet for 20 min, washed 3 times with water and dried overnight before being resuspend in 200 µl methanol and absorbance was measured at 570 nm as described in (Feoktistova et al., 2016). For analysis of cell death with the Incucyte live-cell analysis system (Sartorius), Yoyo3 iodide viability die was used at 1/8000 final concentration and images were acquired. The Incucyte was also used to measure confluency and acquire bright-field images over time with a 10x objective.

### Apoptosis assays

The Annexin V/7AAD kit was used to detect apoptotic cells by flow cytometry according to the manufacturer’s protocol. Caspase 3/7 Glo (Promega) was used to measure caspase 3/7 cleavage. For live cell imaging, BMDMs were cultured in a glass chamber coated with Poly-L-lysine at 37°C and 5% CO_2_. Culture media were supplemented with 2.5mM CaCl2, 20 mM HEPES, propidium iodide (3 µl/well) and Annexin V AF488 (1 µl/well). Images were acquired with a Delta Vision microscope with 10x objective lens every 10 minutes for 24 hours.

### Cell cycle analysis

BMDMs were seeded at 10^6^ cells/well in 6-well low attachment plates and were incubated with 10 µM BrdU for 30 min. In pulse chase experiments, cells were incubated with 10 µM BrdU for 15 min, the media was then replaced, and cells were exposed to dG. At appropriate time points, cells were washed and fixed in 70% ethanol and frozen at -20°C. Cells were washed and resuspend in pepsin solution (1 mg/ml in 30 mM HCl) for 30 min at 37°C, spun down and resuspend in 2M HCl for 15 min at room temperature (RT) and washed with PBS. Cells were then blocked with 0.5% BSA, 0.5% Tween-20 in PBS for 30 min at room temperature, washed and resuspend in FACS buffer with *α*-BrdU AF488 antibody at 1:100 dilution for 30 min at room temperature. Fix Cycle PI/RNAse A staining solution was added to the cells for 30 min at RT. Cells were acquired on a BD LSR II flow cytometer.

### Clonogenic assay

Jurkat cells were treated with dG for 20 hours in IMDM with 10% FCS. 1200 cells were then seeded per well in 6 well plates in methylcellulose, semi-solid medium (40% MethoCult, 39% IMDM, 20% FCS, 1% glutamax and primocin at 100 µg/ml) containing dG. After 13 days incubation, cell colonies were counted manually.

### Measurement of ROS production

H_2_O_2_ production was measured using the ROS-Glo H_2_O_2_ assay (Promega) according the manufacturer’s instructions.

### Western blots

Cells were lysed in NP-40 buffer (150 mM NaCl, 1% NP-40, 50 mM Tris pH 8.0) with protease and phosphatase inhibitors. After 20 min incubation on ice, lysates were centrifuged at 17,000g for 10 min at 4°C. Supernatant was collected and diluted with sample buffer before denaturation at 94°C for 5 min. Samples were loaded on pre-cast 4-12 % gradient Bis-Tris protein gels that were run with MOPS or MES buffer at 120 volts for 2 hours. Transfer to nitrocellulose membranes was performed in transfer buffer (25mM Tris, 192mM glycine, 20% methanol) at 30 volts for 3.5 hours. Membranes were blocked in 5% milk powder in Tris buffered saline with 1 % Tween-20 (TBST) for 1 hour at room temperature then washed 5 times for 5 min in TBST. Membranes were incubated with primary antibody in 5 % milk TBST overnight at 4°C, then washed 5 times for 5 min in TBST. ECL or ECL prime substrates were used for signal detection. In some experiments, membranes were stripped (0.2 M glycine, 0.1 % SDS at pH 2.5) for 15 minutes, washed, blocked and re-probed with a different antibody.

### Retroviral vectors

VSV-G-pseudotyped retroviral vectors were produced by plasmid transfection of 293T cells (Bridgeman et al., 2015). Retroviral infections were performed in the presence of 8 µg/ml polybrene. Bone marrow cells were transduced three times by spin-infection (2500 rpm, 120 min, 32°C, no brakes) on days 1, 2 and 3 of the 7-day differentiation process with the retroviral vector expressing SAMHD1 or a control vector. Cells were seeded into new plates on day 7 and treated with dG on the next day. THP1 cells were transduced by spin infection (2500 rpm, 120 min, 32°C, no brakes), seeded and treated as indicated in the figure legends. Jurkat cells, MDA-MB231 and Hela cells were transduced by adding viral vectors to the culture medium. VLPvpx and VLP_ctrl_ were generated using the SIVmac gag-pol expression vectors SIV3+ and SIV4+ (Maelfait et al., 2016). Human SAMHD1 expression constructs pCSHAwtW and pCSHAk11aW were a kind gift from T. Schaller. These were used to generate lentiviruses for reconstitution of Jurkat cells (Bridgeman et al., 2015). Mouse SAMHD1 expression construct pMSCVpuro-mSAMHD1 was used to generate a retroviral vector to transduce bone marrow cells as described earlier (Rehwinkel et al., 2013).

### Stimulation, staining and mass cytometry analysis of patient samples

PBMCs were collected 24 hours after treatment with forodesine and dG, and were processed according to the Fluidigm Maxpar protocol, using Maxpar reagents. Antibodies are listed in the Supplementary table 2. Cells were collected in 15 ml falcon tubes and were washed in PBS, using centrifugation at 300 g for 5 min. Cells were resuspend at 10^7^/ml in R0 with Cisplatin (1:10,000) and incubated at 37°C for 5 minutes. Cells were washed with R10 and resuspend in Maxpar PBS. Staining was performed on 3*10^6^ cells/tube. Staining with CD56, CD27, CCR4 and CCR7 was done before fixation. Cells were fixed with Maxpar Fix I Buffer at room temperature (RT) for 10 min then washed with Maxpar Cell Staining Buffer (CSB) and spun at 800 g for 5 min. Cells were barcoded (fluidigm barcoding kit) for 30 min at RT, washed twice in CSB, pooled and counted. All further steps were performed on the pooled sample. Cells were blocked in FcR block diluted in CSB (1:10) for 10 min at RT. Surface staining antibody mix was added to blocking solution and incubated for 30 min at RT. Cells were washed in CSB, resuspend in ice cold methanol added dropwise under the vortex and stored at -80°C overnight. Cells were washed twice with CSB and stained with the intracellular antibody mix for 30 min at RT and stained with intercalator overnight. Next day they were washed with CSB and resuspend in water before acquisition on the Helios mass cytometer (Fluidigm). Samples were normalized, concatenated and de-barcoded using Helios software. Files were analysed with Cytobank.

### Generation of *Samhd1^-/-^* B16F10 cells

sgRNAs were designed to excise exon 2 of mouse *Samhd1* (Gene ID: 56045). Exon 2 is critical to both isoforms of *Samhd1* (see Supplementary figure 2b and (Rehwinkel et al., 2013)). B16F10 cells were co-transfected with pX458-Ruby-sgRNA-1 and pX458-sgRNA-2 plasmids. GFP-Ruby double positive cells were single cell sorted and clones were expanded. A PCR screening approach was used to identify knock-out cells. PCR-1 was designed to amplify a long fragment (709 bp) from the WT allele and a short fragment (350 bp) from the KO allele using primer 1 fwd and primer 2 rvs (see Supplementary figure 2b). PCR-2 had a primer located in exon 2 and amplified a fragment (352 bp) only from the WT allele using primer 2 rvs and primer 3 fwd (Supplementary table 1).

### Statistics

All experiments were performed three times or more independently under similar conditions, unless specified otherwise in figure legends. Statistical significance was calculated by unpaired t-test, one-way ANOVA or two-way ANOVA as described in the figure legends; p<0.05 was considered significant. Graph pad prism 7 software was used to generate graphs and to perform statistical analysis.

### Study approval

Mouse work was performed in accordance with the UK Animals (Scientific Procedures) Act 1986 and institutional guidelines for animal care. This work was approved by a project license granted by the UK Home Office (PPL No. PC041D0AB) and also was approved by the Institutional Animal Ethics Committee Review Board at the University of Oxford.

Informed consent from all patients was obtained in line with the Declaration of Helsinki. The CLL work was covered by the Ethics approval REC 09/H1306/54. Human fibroblasts from patients with AGS were collected with approval by a U.K. Multicentre Research Ethics Committee (reference number 04:MRE00/19).

## Author contributions (using <u>the CRediT taxonomy</u>)

Conceptualization: T.D., R.E.R., A.S. and J.R.; Methodology: T.D., R.E.R., C.C. and A.B.; Software: n.a.; Validation: T.D. and J.R.; Formal analysis: T.D. and J.R.; Investigation: T.D., B.D., R.E.R. and S.S.; Resources: J.K., P.H. and A.S.; Data curation: T.D.; Writing – Original Draft: T.D. and J.R.; Writing – Review & Editing: all authors; Visualization: T.D. and J.R.; Supervision: J.R., A.C. and K.D.K; Project administration: J.R.; Funding acquisition: T.D. and J.R.

## Acknowledgments

The authors thank Y. Crow and G.I. Rice for providing primary human fibroblasts, T. Yokomatsu for homo-DFPP-DG or 6C-DFPP-DG, and T. Schaller for SAMHD1 expression vectors. We thank Quentin Sattentau, members of the Rehwinkel lab, Jonathan Maelfait, Persephone Borrow, Jane McKeating, Philippe Pasero, Yea-Lih Lin, Skirmantas Kriaucionis, Wojciech Niedzwiedz and Vincenzo Cerundolo for discussion. We would like to acknowledge Giorgio Napolitani and Michalina Mazurczyk for their help in the mass cytometry facility at the WIMM for providing technical expertise, cell analysis services and scientific input. The facility is supported by the MRC HIU core funded project, reference MC_UU_00008, and the Oxford Single Cell Biology Consortium (OSCBC). We thank Philip Hublitz for his help with generation of *Samhd1^-/-^* B16F10 cells. The WIMM Genome Engineering Facility is supported by grants from the MRC/MHU (MC_UU 12009), the John Fell Fund (123/737) and by WIMM Strategic Alliance awards G0902418 and MC_UU_12025. The authors would like to acknowledge Dr Melani Oates, Liverpool Bio-Innovation Hub Biobank, for help with CLL sample retrieval. This work was funded by the UK Medical Research Council [MRC core funding of the MRC Human Immunology Unit; J.R.]; the Wellcome Trust [grant number 100954; J.R.], the Swedish Cancer Society and the Swedish Research Council [A.C.] and a C1 KU Leuven Research Council grant [number C14/18/104; KDK]. T.D. was supported by the Wellcome Trust Infection, Immunology & Translational Medicine doctoral programme [grant number 105400/Z/14/Z]. AS is partly funded by the National Institute for Health Research Oxford Biomedical Research Centre. The views and opinions expressed are those of the authors and do not necessarily reflect those of the National Institute for Health Research, the UK National Health Service, the UK Department of Health or the Universities of Oxford and Cambridge. The funders had no role in study design, data collection and analysis, decision to publish, or preparation of the manuscript.

## Data availability statement

The authors declare that all data supporting the findings of this study are available within the paper and its supplementary information files.

## Supplementary Information

### Supplementary figures

**Supplementary Figure 1.**
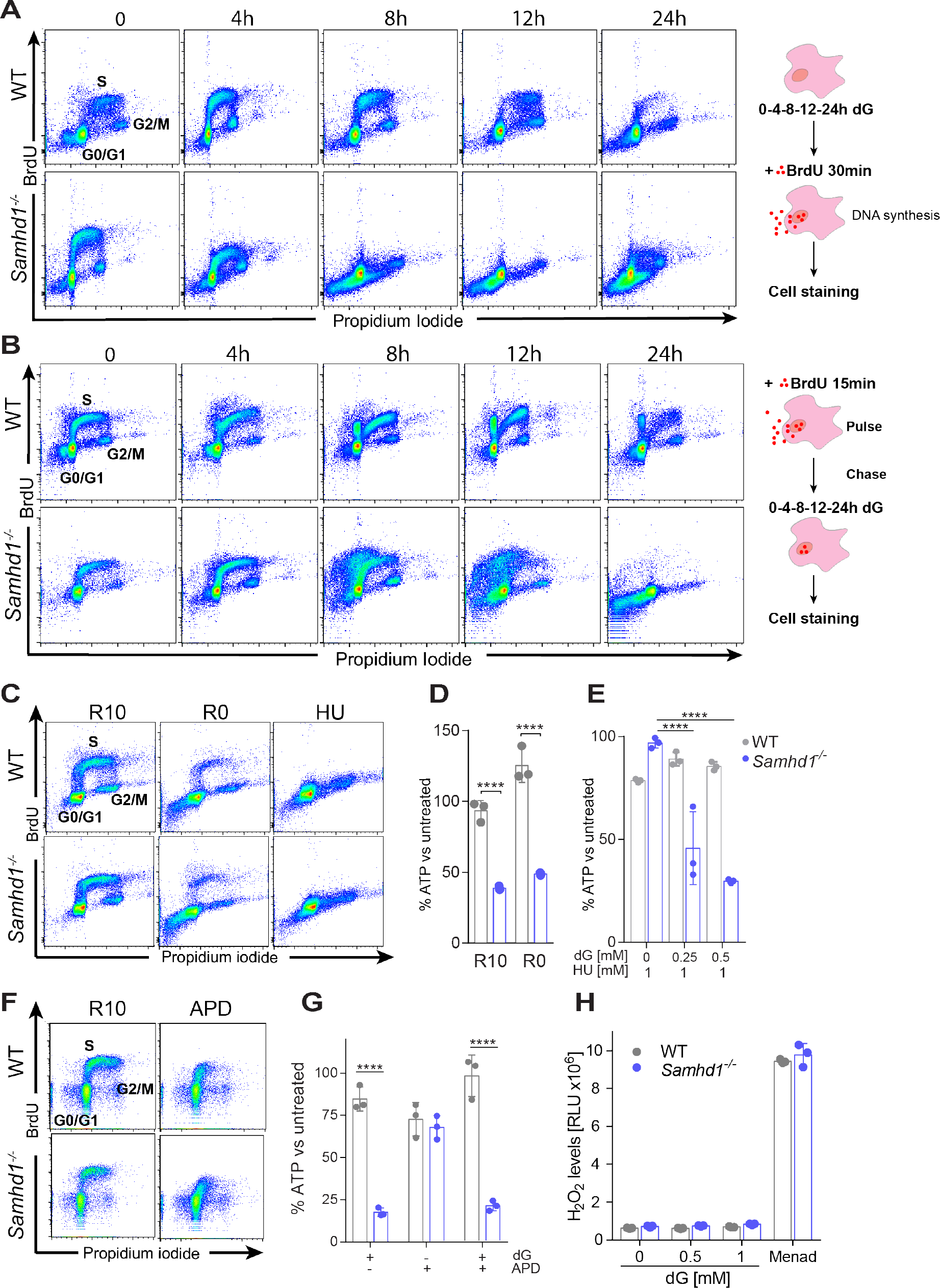
dG-induced cell death in *Samhd1*^-/-^ cells is independent of nuclear DNA replication, related to Figure 2. (A) BMDMs were treated with 0.5 mM dG for the indicated periods of time. Cells were then labelled with BrdU for 30 minutes and fixed. Cells were stained using *α*-BrdU antibody and PI and analysed by flow cytometry. (B) BMDMs were labelled with BrdU for 15 minutes and then treated with 0.5 mM dG for the indicated periods of time. Cells were then analysed as in (**A**). (**C-E**) After 7 days of conventional culture, BMDMs were grown in medium containing 10% FCS (R10) or in serum-free medium (R0) for 24 hours. Alternatively, BMDMs were treated with 1 mM hydroxyurea (HU) for 8 hours. (**C**) Cells were analysed as in (**A**). (**D**) Cells were treated with 0.5 mM dG for 24 hours and viability was analysed as described in Figure 1A. (**E**) Cells were treated with 0.5 mM dG for 16 hours and viability was analysed as described in Figure 1A. (**F-G**) BMDMs were treated or not with 1 µM of aphidicolin (APD) in R10 for 16 hours. (**F**) Cells were analysed as in (**A**). (**G**) Cells were treated with 0.5 mM dG for 24 hours and viability was analysed as described in Figure 1A. (**H**) H_2_O_2_ production was measured using the ROS-Glo H_2_O_2_ assay (Promega). BMDMs were treated with the indicated concentrations of dG for 3 hours. The H_2_O_2_ substrate solution was then added for 6 hours. Control cells were treated with 20 µM Menadione (Menad) during this period. Luminescence was measured after addition of the detection solution. Data are representative of three (**A-G**) or two (**H**) independent experiments, respectively. In panels **D-E** and **G-H** data from triplicate measurements are shown with mean ± SD. P-values determined with two-way ANOVA are indicated. **** p<0.0001.

**Supplementary Figure 2.**
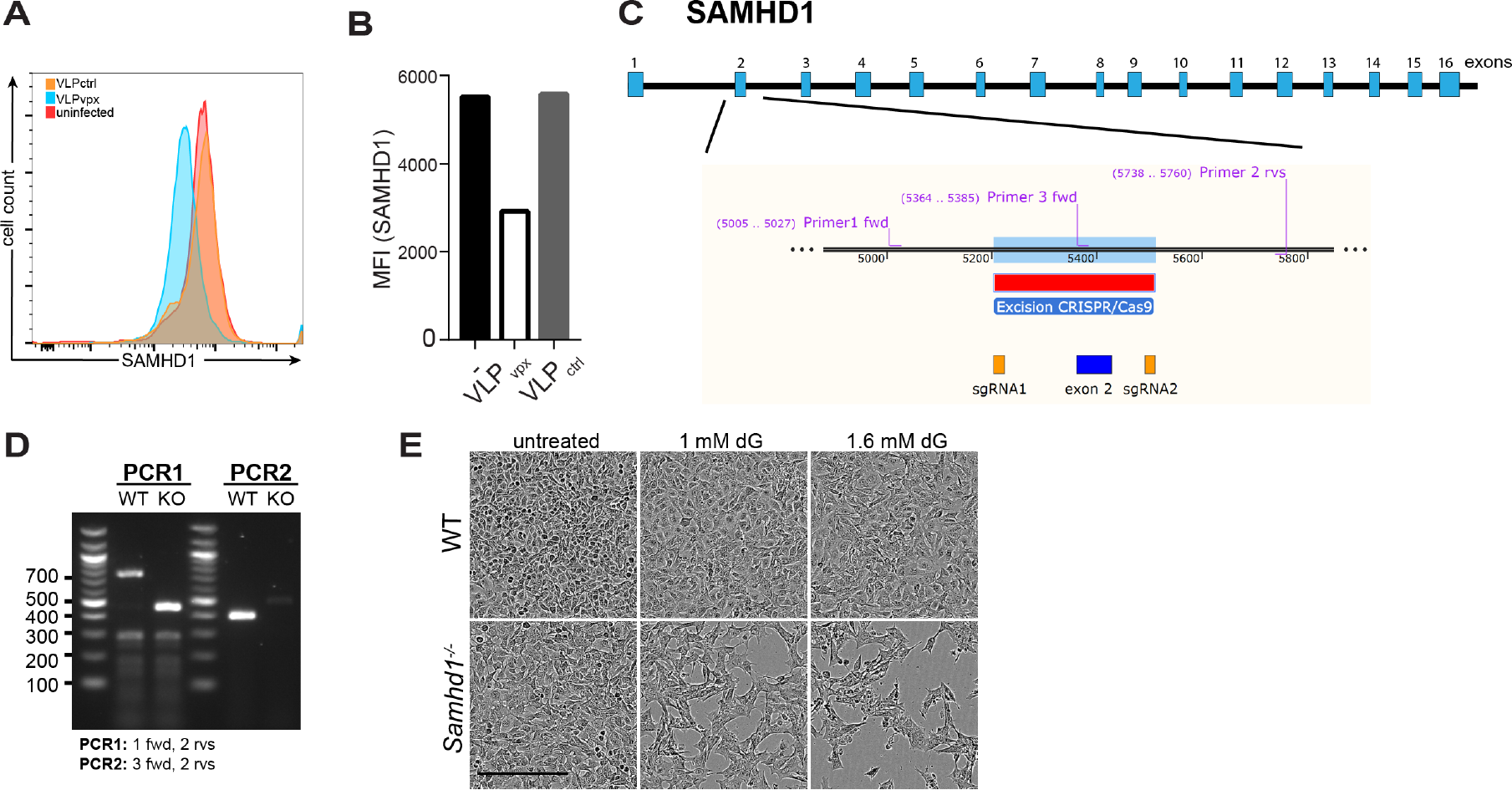
Validation of SAMHD1-deficient cells, related to Figure 3. (**A,B**) HeLa treated as described in Figure 3A,B were stained with *α*-SAMHD1 antibody and analysed by flow cytometry. Data are represented as histograms (**A**) and the mean fluorescence intensity (MFI) of the SAMHD1 signal is shown in (**B**). (**C,D**) Representation of the CRISPR/Cas9 strategy to generate B16F10 *Samhd1^-/^*^-^ cells used in Figure 3E-G. The knock-out of *Samhd1* exon 2 was validated by PCR using the indicated primers (**D**). (**E**) Brightfield images of WT and *Samhd1^-/^*^-^ B16F10 cells 20 hours after treatment with dG from the experiment in Figure 3G are shown. The scale bar represents 300 µm. Panels **A,B** and **E** are representative of three independent experiments, respectively.

**Supplementary Figure 3.**
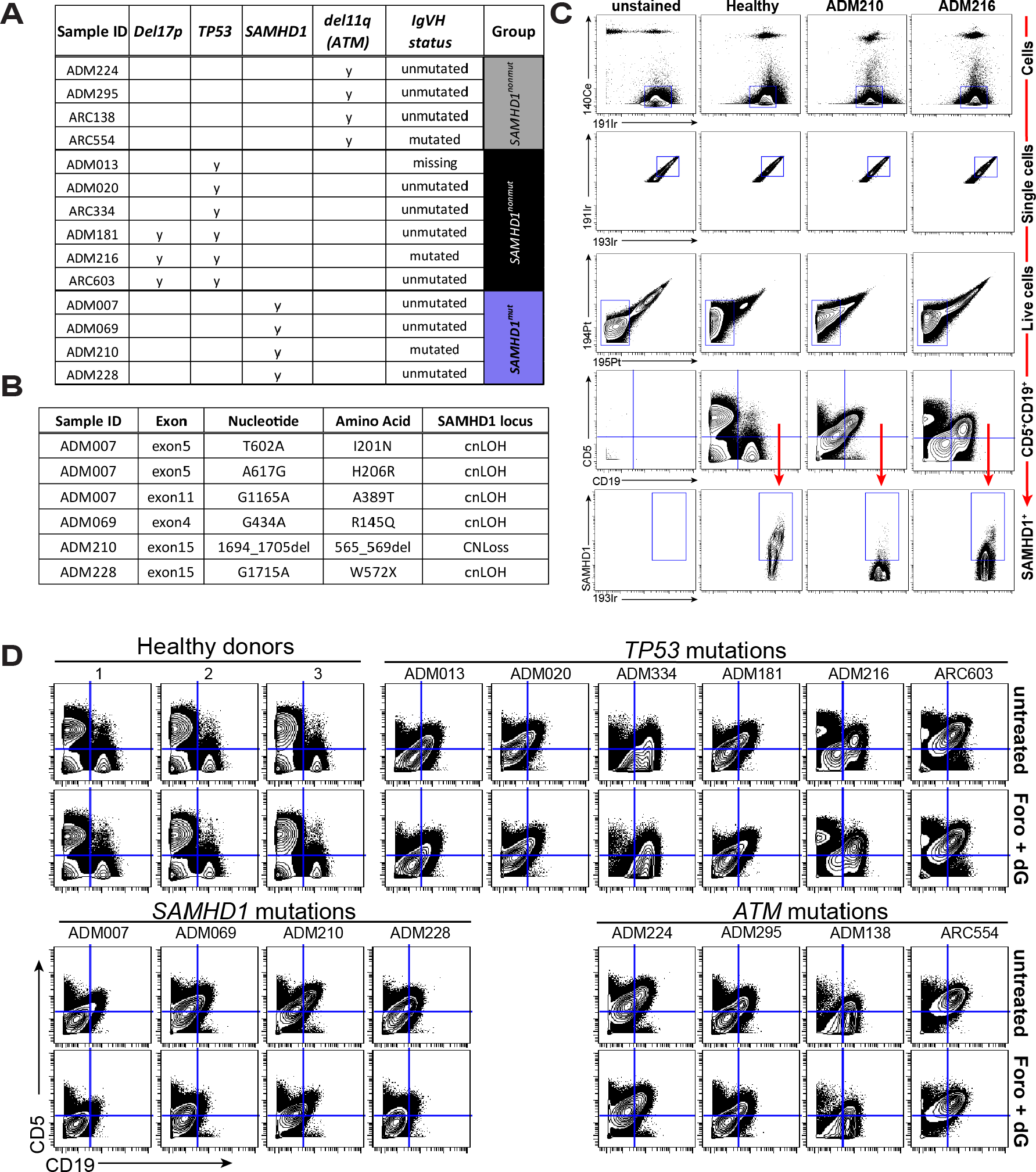
CyTOF analysis, related to Figure 5. (A) List of CLL samples and mutation status. CLL samples carrying either *TP53* lesions (Del17p or mutation) or *ATM* lesions (Del11q) served as controls for *SAMHD1* mutated samples. IgVH, immunoglobulin variable heavy chain gene. (B) Details of *SAMHD1* mutations. cnLOH, copy neutral loss of heterozygosity; CNLoss, copy number loss. (C) CyTOF gating strategy for the experiment shown in Figure 5. In brief, the 140Ce channel was used to exclude calibration beads. The 191Ir and 193Ir channels detect cells labelled with an intercalator and were used to identify single cells. The 194Pt and 195Pt channels were used to remove dead (cisplatin-labelled) cells from subsequent analysis of CD5, CD19 and SAMHD1 expression. (D) Expression of CD5 and CD19 is shown as in Figure 5B for all analysed samples.

**Supplementary Figure 4.**
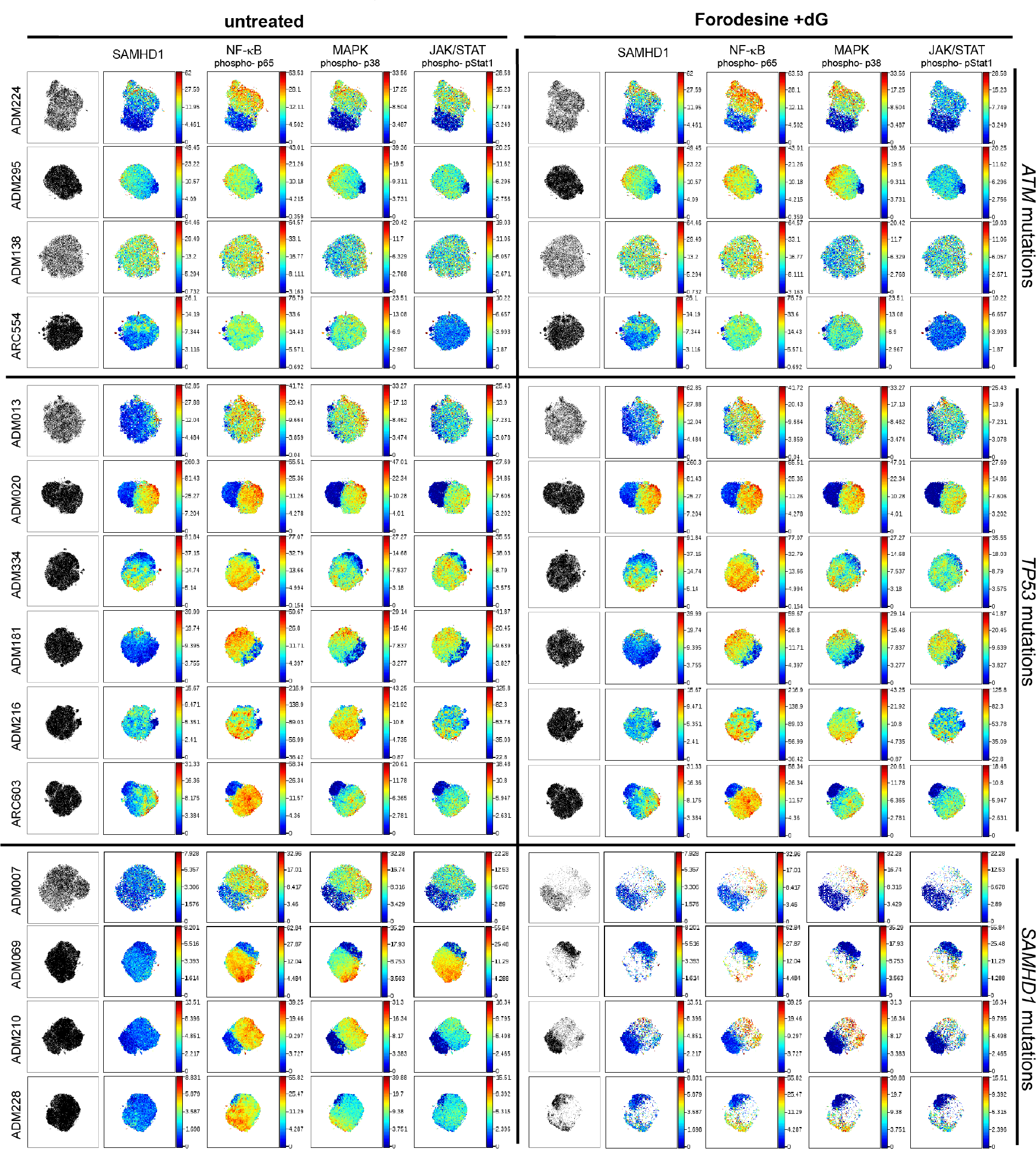
CLL patient cells with *SAMHD1* mutation are more sensitive to Forodesine and dG treatment, related to Figure 5. CyTOF data from each sample were analysed separately by viSNE analysis as described in Figure 5E. Data from all patients studied are shown; for completeness, we included here those selected for Figure 5E.

**Supplementary Figure 5.**
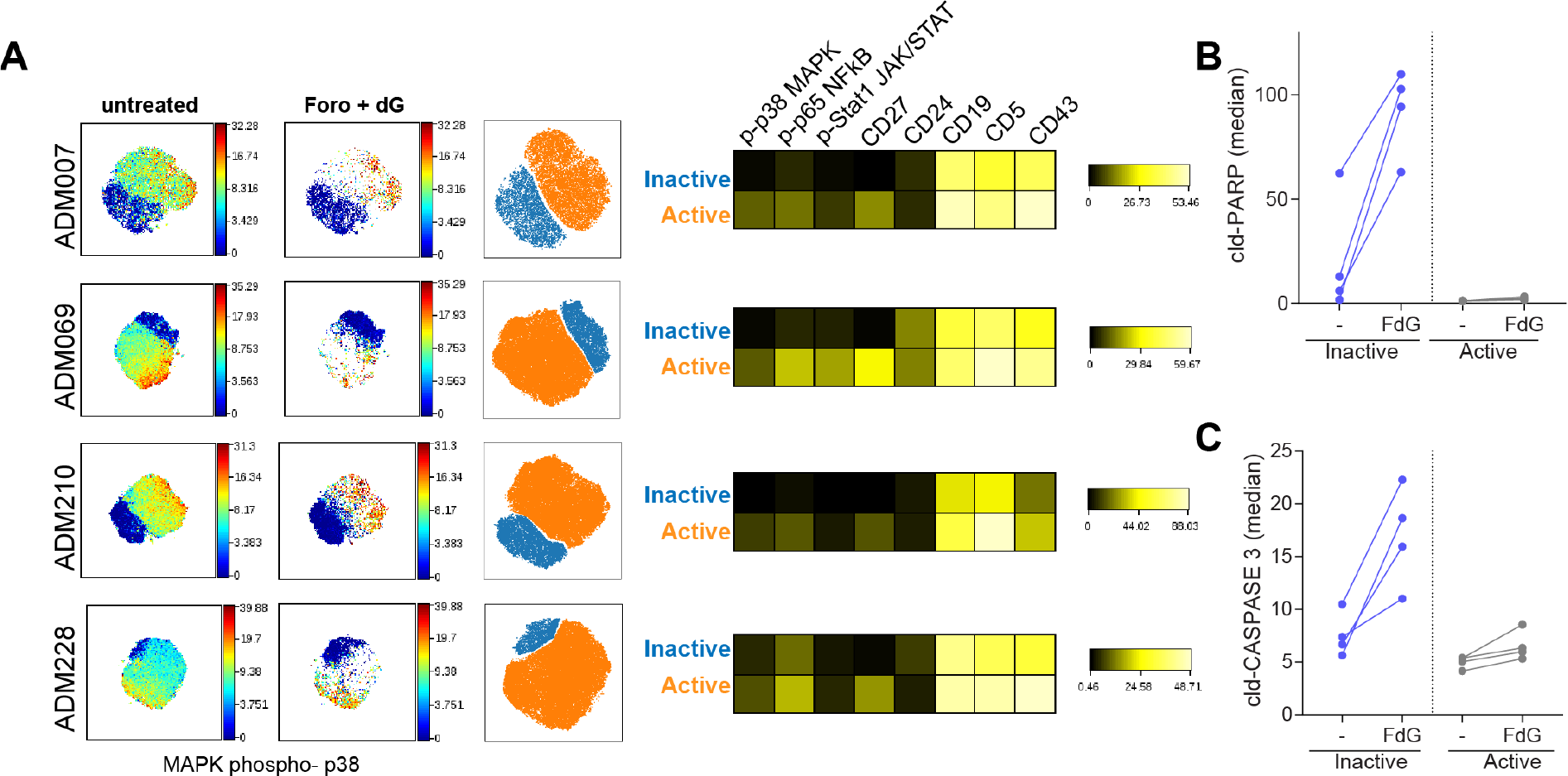
Forodesine and dG induce apoptosis in “inactive” CLL B-cells, related to Figure 5. (**A**) tSNE plots using live CD5^+^CD19^+^ cells from *SAMHD1* mutated patients, coloured by p-p38 levels, were used to manually gate “inactive” [blue] and “active” [orange] cells (left). Expression of selected markers in these sub-populations is shown as a heat map (right). (**B,C**) The staining for cleaved-PARP (**B**) and cleaved-CASPASE3 (**B**) was analysed in “inactive” and “active” cells. Median values are shown in untreated and treated cells. FdG, treatment with 2 µM forodesine and 20 µM dG.

### Supplementary tables

**Supplementary Table 1.**
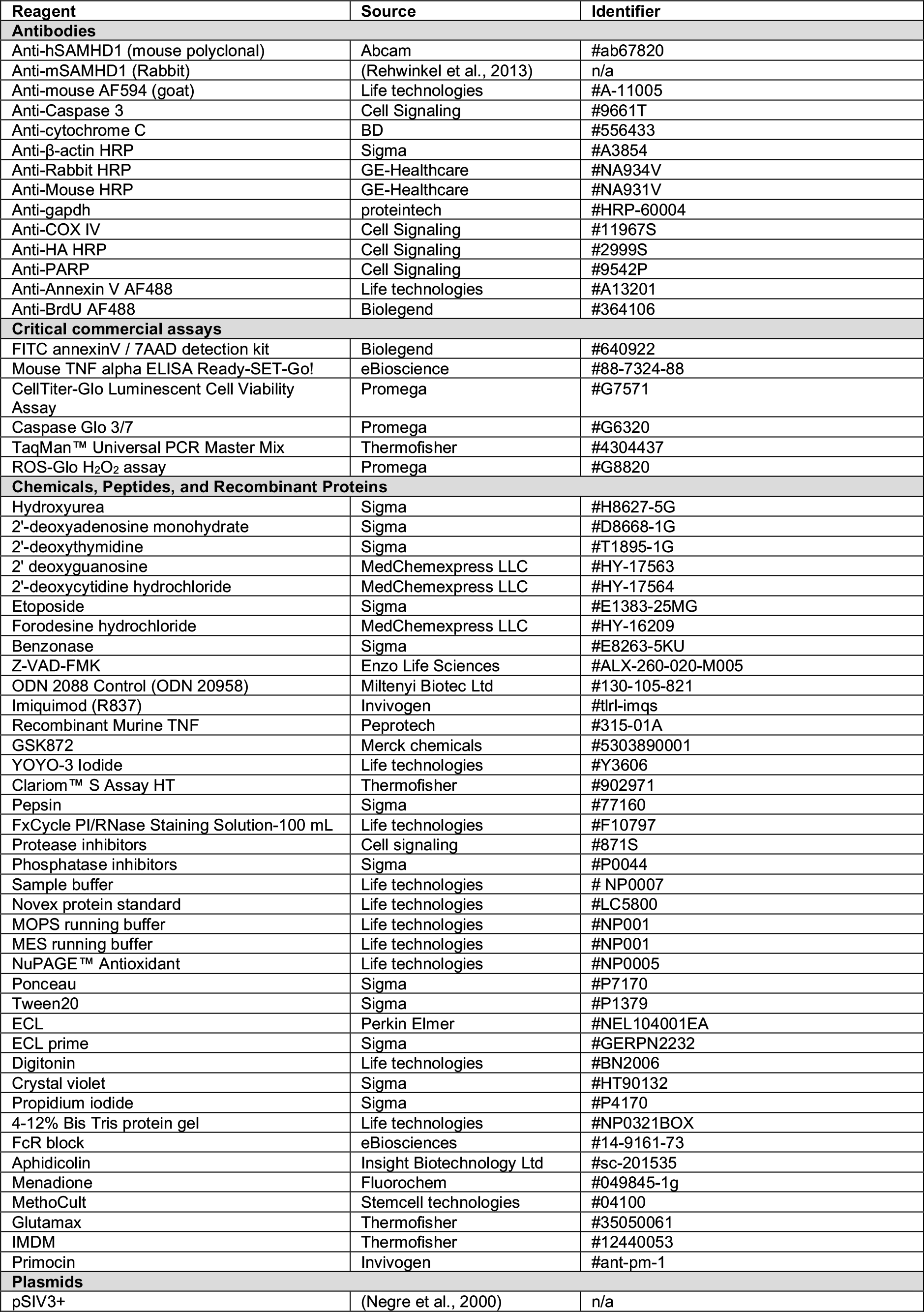
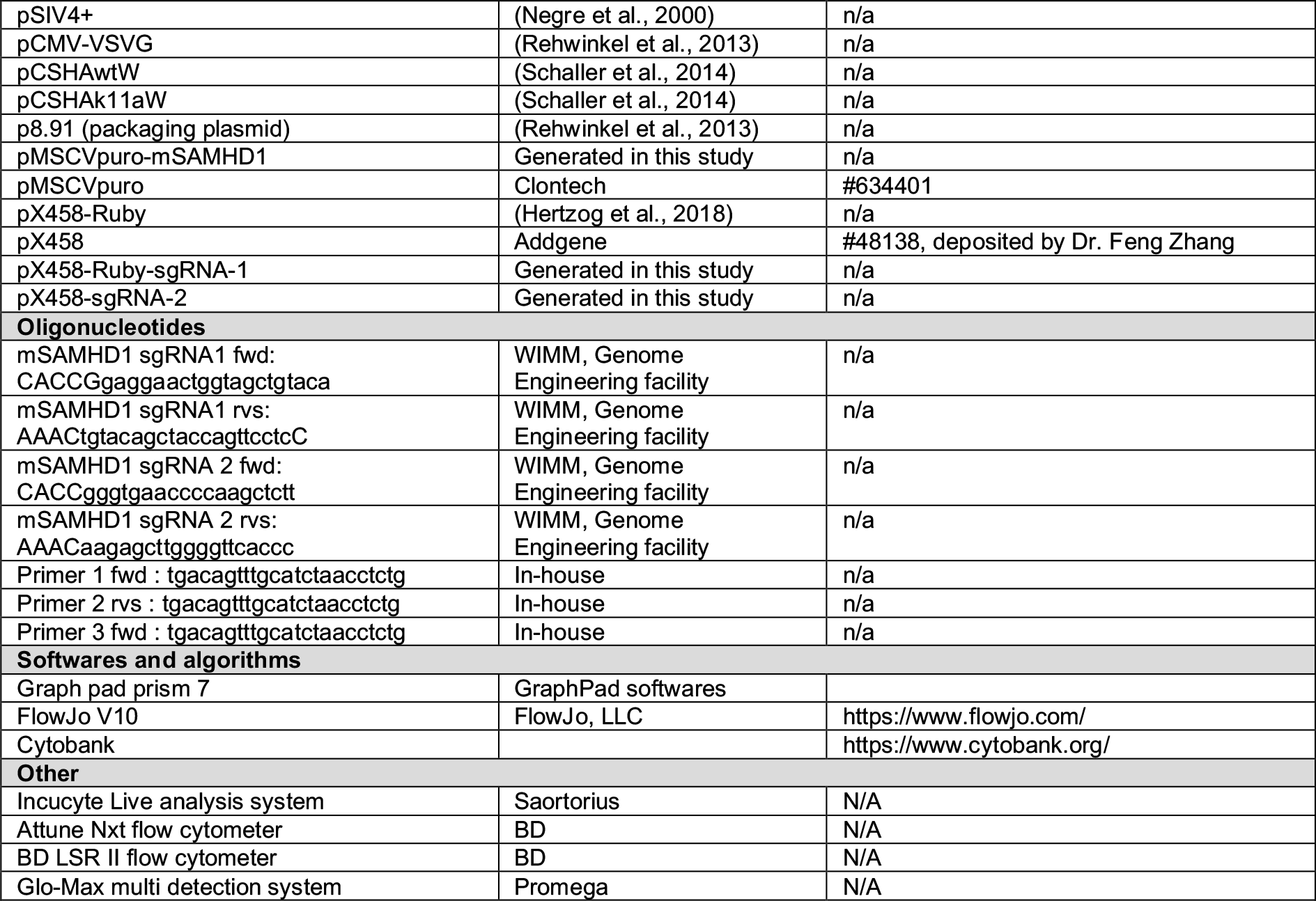
Reagents

**Supplementary Table 2.** CyTOF antibody panel.

| Target | Clone | Company | Catalog | Location |
| --- | --- | --- | --- | --- |
| CD45 | HI30 | Fluidigm | 3089003B | surface |
| CD11c | Bu15 | Biologend | 337221 | surface |
| CD11b (Mac-1) | ICRF44 | Biologend | 301337 | surface |
| CD4 | RPA-T4 | Fluidigm | 3145001B | surface |
| CD20 | 2H7 | Fluidigm | 3147001B | surface |
| CD56 (NCAM) | NCAM16.2 | Fluidigm | 3149021B | surface |
| CD123 | 6H6 | Biologend | 306027 | surface |
| CD27 | O323 | Biologend | 302839 | surface |
| CD38 | HIT2 | Biologend | 303535 | surface |
| CD45RA | HI-100 | Biologend | 304143 | surface |
| CD14 | M5E2 | Fluidigm | 3160001B | surface |
| CD23 | EBVCS-5 | Fluidigm | 3164018B | surface |
| CD197 (CCR7) | G043H7 | Fluidigm | 3167009A | surface |
| CD8a | SK1 | Fluidigm | 3168002B | surface |
| CD24 | ML5 | Fluidigm | 3169004B | surface |
| CD3 | UCHT1 | Fluidigm | 3170001B | surface |
| CD19 | HIB19 | Biologend | 302247 | surface |
| CD5 | UCHT2 | Biologend | 300627 | surface |
| HLA-DR | L243 | Fluidigm | 3174001B | surface |
| CD194 (CCR4) | 205410 | Fluidigm | 3175021A | surface |
| CD43 | eBio84-3C1 | Invitrogen | 14-0439-82 | surface |
| CD16 | 3G8 | Fluidigm | 3209002B | surface |
| cld-CASP3 | 5AE1 | Fluidigm | 3172023A | intracellular |
| cld-PARP | F21-852 | Fluidigm | 3143011C | intracellular |
| pStat1 [Y701] | 4a | Fluidigm | 3153005A | intracellular |
| p38 [T180/Y182] | D3F9 | Fluidigm | 3156002A | intracellular |
| SAMHD1 | polyclonal | Abcam | Ab67820 | intracellular |
| pNFκBp65 [S529] | K10x | Fluidigm | 3166006A | intracellular |

